# IL-12/18/21 pre-activation enhances the anti-tumor efficacy of expanded γδT cells and overcomes resistance to anti-PD-L1 treatment

**DOI:** 10.1101/2022.08.11.503563

**Authors:** Huey Yee Teo, Yuan Song, Kylie Su Mei Yong, Yonghao Liu, Yu Mei, Zuhairah Binte Hanafi, Ying Zhu, Nicholas R. J. Gascoigne, Qingfeng Chen, Haiyan Liu

## Abstract

γδT cells are promising candidates for cellular immunotherapy due to their immune regulation through cytokine production and MHC-independent direct cytotoxicity against a broad spectrum of tumors. However, current γδT cell-based cancer immunotherapy has limited efficacies and novel strategies are needed to improve its clinical outcomes. Here, we report that cytokine pre-treatment with IL-12/18, IL-12/15/18, IL-12/18/21 and IL-12/15/18/21 could effectively enhance the activation and cytotoxicity of *in vitro* expanded murine and human γδT cells. However, only adoptive transfer of IL-12/18/21 pre-activated γδT cells significantly inhibited tumor growth in a murine melanoma model and a hepatocellular carcinoma model. Both IL-12/18/21 pre-activated antibody-expanded and zoledronate-expanded human γδT cells effectively controlled tumor growth in a humanized mouse model. IL-12/18/21 pre-activation promoted γδT cell proliferation and cytokine production *in vivo* and enhanced IFN-γ and TNF-α production, as well as granzyme B expression by endogenous CD8^+^ T cells in a cell-cell contact dependent manner. Furthermore, the adoptive transfer of IL-12/18/21 pre-activated γδT cells could overcome the resistance to anti-PD-L1 therapy, and the combination therapy had a synergistic effect on the therapeutic outcomes. Moreover, the enhanced anti-tumor function of adoptively transferred IL-12/18/21 pre-activated γδT cells was largely diminished in the absent of endogenous CD8^+^T cells when administered alone or in combination with anti-PD-L1, suggesting a CD8^+^T cell-dependent mechanism. Taken together, IL-12/18/21 pre-activation could promote γδT cell anti-tumor function and overcome the resistance to checkpoint blockade therapy, indicating an effective combinational cancer immunotherapeutic strategy.

**Synopsis:**

1. IL-12/18/21 pre-activation enhances the anti-tumor efficacy of adoptively transferred γδT cells by promoting its proliferation, activation, and cytotoxicity, as well as activating endogenous CD8^+^ T cells.
2. Adoptive transfer of IL-12/18/21 pre-activated γδT cells could overcome the resistance to anti-PD-L1 therapy, indicating an effective combinational cancer immunotherapeutic strategy.

## Introduction

Cancer adoptive immunotherapy is the delivery of immune effector cells that are gene-modified or expanded *in vitro* to patients to fight cancer. Chimeric antigen receptor T (CAR-T) cell adoptive immunotherapy has provided beneficial clinical outcome in hematological malignancies while its therapeutic efficacy in solid tumors remains a challenge(1). Several clinical studies have shown that γδT cells display efficient anti-tumor function in a range of cancers, from renal cell carcinoma, lung cancer to hematological malignancies(2-4). There are numerous advantages that γδT cells could offer compared to αβT cells thus they are considered to be a promising tool in cancer immunotherapy. Unlike αβT cells that require major histocompatibility complex (MHC) class I and II presentation, γδT cells respond to stress-induced ligands such as MIC-A, MIC-B and ULBP6 and thus are not MHC restricted(5). Because of this feature, γδT cells are able to recognize and distinguish tumor cells from normal cells through the accumulation of phosphorylated antigens, known as phosphoantigens, which are produced by the tumor cells due to higher metabolism and usage of the mevalonate isoprenoid pathway(6,7). Both *in vitro* and *in vivo* studies have demonstrated that γδT cells are able to recognize a range of tumor cells and exert anti-tumor capabilities(7). The ability to recognize stress-induced ligands also allows fast response from γδT cells in comparison to αβT cells which require antigen processing and presentation via antigen presenting cells(8). Besides its innate cytotoxic ability, γδT cells also have antigen-presenting capability upon stimulation(9). γδT cells could eradicate tumor cells and present tumor-associated antigens to αβT cells(8), leading to a long-lasting and dynamic immune response. Being able to release abundant pro-inflammatory cytokines such as interferon-gamma (IFN-γ) and tumor necrosis factor-alpha (TNF-α) upon stimulation is another important characteristic of γδT cells(10-12). IFN-γ-producing γδT cells are able to upregulate MHC class I expression by tumor cells, leading to the activation of CD8^+^ T cell response which enhances tumor clearance(13). Therefore, these unique features of γδT cells make them favorable effector cells in adoptive cancer immunotherapy.

Many recent studies have demonstrated the potent activity of γδT cells against various solid tumors and leukemia(14-17), however, not all γδT cell immunotherapy exerted effective anti-tumor efficacy(18-20). The adoptively transferred γδT cells can become dysfunctional and exhausted in the tumor microenvironment (TME) probably due to the presence of suppressive immune subsets or hypoxic environment(21). It was also found that γδT cells disappeared from circulation over time in clinical trials(22). The survival and proliferation of adoptively transferred γδT cells *in vivo* limited the success of γδT cells therapy. Moreover, the pro-tumorigenic activity of intra-tumoral γδT cells have been reported(23,24). IL-17-producing γδT cells could recruit myeloid-derived suppressor cells (MDSCs) in the TME to inhibit anti-tumor T cell response(25). Therefore, novel strategies are needed to differentiate and generate long-lived anti-tumor γδT cells to boost the therapeutic efficacy of γδT cell-based therapy.

T cell functions are known to be enhanced by various cytokines. Interleukin (IL)-12 is a pro-inflammatory cytokine which can promote the cytotoxic activity of NK cells and T cells by inducing IFN-γ production(26). IL-12 interacts synergistically with IL-18 to boost the cytotoxicity of Th1 cells by reciprocally upregulating each other’s receptor expression(27-29). IL-18 can independently enhance perforin-dependent cytotoxicity and FasL-mediated cytotoxicity in NK cells(30,31). Moreover, IL-15 and IL-21 are able to synergize and induce proliferation and IFN-γ production in CD8^+^ T cells *in vitro*(32). One interesting feature of IL-21 is that IL-21-primed CD8^+^ T cells have memory phenotypes and can better persist *in vivo(33)*. This would be advantageous in generating a durable and long-term response against tumors. Although the functions of these cytokines on αβT cells and NK cells are relatively well-studied, their functions, especially their combinatory effects on γδT cells are not clear. In *ex vivo* expansion settings, IL-15 was able to boost proliferation, strengthen the polarization towards a Th1 phenotype and enhance the cytotoxic capability in Vγ9Vδ2 T cells against several tumor cells(34). The effect of IL-21 on γδT cells is similar to IL-15(35). IL-18 could potently expand γδT cells with effector memory phenotypes and anti-tumor activity(36). A combination of IL-12/IL-18 increased levels of granzymes and IFN-γ together with decreased expression levels in co-inhibitory receptor, BTLA, on Vγ9Vδ2 T cells(37,38). Since systemic administration of cytokines would cause severe undesirable side effects(39), pre-activation of immune cells prior to adoptive transfer is a safer and more targeted delivery of the anti-tumor effect of these cytokines. This approach has been demonstrated on NK cells, where combined activation with IL-12, IL-15 and IL-18 prolonged the NK cell survival and enhanced its effector function(40).

Immune checkpoint blockade immunotherapy has led to important clinical advances via regulating the T cell immune responses. It showed unprecedented durable responses in recent clinical trials with a variety of cancers(41). However, immune checkpoint blockade is limited by primary and acquired resistance mechanism, including the lower expression of PD-L1 in tumors and the lack of T cell infiltration in the TME. Combination therapy which aims to increase the infiltration of T cells into the TME or change the gene expression profile of tumor cells is a great strategy to optimize the checkpoint blockade immunotherapy. Thus, combining with γδT cell-based adoptive therapy which provides IFN-γ and infiltrating T cells may overcome the resistance to checkpoint blockade immunotherapy.

In the current study, we aim to improve the efficacy of γδT cell-based therapy by pre-activation with a combination of cytokines. The combinations of IL-12/18, IL-12/15/18, IL-12/18/21, and IL-12/15/18/21 significantly promoted the expressions of CD25, FasL, perforin, and the production of IFN-γ and TNF-α. The killing capacity of murine γδT cells against tumor cells was also promoted by the cytokine pre-activation. Similar results were obtained with human γδT cells. To further evaluate the anti-tumor effect of the cytokine pre-activated γδT cells *in vivo*, we adoptively transferred IL-12/18, IL-12/15/18 and IL-12/18/21 pre-activated γδT cells in the murine subcutaneous melanoma and orthotopic hepatocellular carcinoma (HCC) model. Although all cytokine pre-activated γδT cells exhibited enhanced proliferation and survival, only IL-12/18/21 combination significantly inhibited tumor development in both models. IL-12/18/21 pre-activated γδT cells maintained higher IFN-γ production *in vivo* and significantly promoted the production of granzyme B, IFN-γ and TNF-α by endogenous CD8^+^ T cells. IL-12/18/21 pre-activated human γδT cells also inhibited tumor growth in a humanized mouse model. Moreover, IL-12/18/21 pre-activated γδT cell adoptive therapy had a synergistic anti-tumor effect when combined with anti-PD-L1 treatment, suggesting it could overcome the resistance to checkpoint blockade therapy. Thus, IL-12/18/21 pre-activation could be an effective strategy to enhance the anti-tumor efficacy of adoptively transferred γδT cells and could be combined with immune checkpoint blockade therapy to achieve optimal clinical outcomes.

## Materials and Methods

### Cell lines

A20 murine lymphoma cell line, B16-F10 murine melanoma cell line, Hepa1-6 murine hepatoma cell line, K562 human myelogenous leukemia cell line, and HepG2 human HCC cell line were purchased from American Type Cell Culture (ATCC, Manassas, VA). SMMC-7721 human HCC cell line was purchased from Cell Bank, Chinese Academy of Sciences (Beijing, China). A20 and K562 were cultured with RPMI 1640 while B16-F10, Hepa1-6, HepG2 and SMMC-7721 were cultured with DMEM (Hyclone, Chicago, IL). RPMI 1640 and DMEM were supplemented with 10mM HEPES (Hyclone, Chicago, IL), 1mM sodium pyruvate (Hyclone, Chicago, IL), MEM non-essential amino acids (Life Technologies, Carlsbad, CA), penicillin-streptomycin (Hyclone, Chicago, IL), 10% heat-inactivated fetal bovine serum (Hyclone, Chicago, IL) and beta-mercaptoethanol (Sigma-Aldrich, St. Louis, MO). Cell lines have been recently authenticated and were periodically tested for mycoplasma species and were confirmed negative.

### Mice

C57BL/6 mice (aged 6-8 weeks) were purchased from InVivos (Singapore) and were used for the initial screening of the effects of cytokine combinations on γδT cells and *in vivo* experiments. TCR-β knockout mice (B6.129P2-Tcrbtm1Mom/J) were purchased from The Jackson Laboratory (Bar Harbor, ME) and bred in National University of Singapore animal housing facility. CD45.1 C57BL/6 mice were obtained from Dr. Nicholas R. Gascoigne’s lab at National University of Singapore and were cross-bred with TCR-β knockout mice for the culture of CD45.1 γδT cells. All mice were kept in specific pathogen free facility and all animal experiments have been conducted in accordance with the approval of the National University of Singapore Institutional Animal Care and Use Committee (murine subcutaneous melanoma model: NUS IACUC#R20-1462, murine HCC model: NUS IACUC#R19-1173).

### Expansion of murine γδT cells

6-well plates were coated with anti-TCR γδ antibody at 10µg/ml overnight. Spleen of CD45.1/CD45.2-TCR-β knockout C57BL/6 mouse, which are deficient of αβT cells, were homogenised and filtered. Red blood cells were removed by incubating with ACK solution for 5 minutes. Cells were washed with RPMI 1640 twice and then seeded at a density of 2.5×10^6^ cells/ml. Anti-mouse-CD28 antibody (1µg/ml) (Biolegend, San Diego, CA) and 100IU/ml recombinant human IL-2 (Miltenyi Biotec, Bergisch Gladbach, Germany) were added. A purity of more than 98% γδT cells were confirmed by flow cytometry on day 5.

### Expansion of human γδT cells

Peripheral blood mononuclear cells (PBMCs) from healthy donors’ whole blood or apheresis cone from Health Sciences Authority, Singapore, were obtained. Written informed consent was obtained from donors. This study was conducted in accordance with the approval of the National University of Singapore Institutional Review Board (NUS-IRB-H-17-044 and NUS-IRB-2020-584). PBMCs were isolated with Ficoll-Hypaque (GE Healthcare, Chicago, IL) density gradient centrifugation. Plasma was collected. Erythrocytes were removed with ACK solution for 5-7 minutes. Cells were washed with RPMI 1640 twice and then seeded at a density of 1×10^6^ cells/ml. For Zol stimulation, 5uM of zoledronate (Zol) (Sigma-Aldrich, St. Louis, MO), recombinant human IL-2 (100IU/ml) and 10% plasma from the same donor were added to culture. For antibody stimulation, cells were seeded into pre-coated wells with anti-TCR pan-γδ clone IMMU510 antibody (5ug/ml) (Beckman Coulter, Krefeld, Germany) and IL-2 (100IU/ml). On day 3, cells were expanded accordingly and maintained with IL-2 (100IU/ml). Cells were then checked for the percentages of Vδ1 and Vδ2 subsets between day 8 and 12 by flow cytometry. This study was approved by National University of Singapore-Institutional Review Board.

### Cytokine pre-activation of γδT cells

IL-2 was included in control and all cytokine combinations. The following cytokines were added at the indicated concentrations for murine γδT cells: recombinant murine IL-12 (10ng/ml) (Peprotech, Rocky Hill, NJ), IL-15 (50ng/ml) (Peprotech, Rocky Hill, NJ), IL-18 (100ng/ml) (R&D Systems, Minneapolis, MN) and IL-21 (30ng/ml) (eBioscience, San Diego, CA). For human γδT cells, recombinant human IL-12 (Stemcell Technologies, Vancouver, Canada), IL-15 (Miltenyi Biotec, Bergisch Gladbach, Germany), IL-18 (R&D Systems, Minneapolis, MN) and IL-21 (Miltenyi Biotec, Bergisch Gladbach, Germany) were added at a concentration of 10ng/ml. Cytokines were added on day 5-6 to murine γδT cells and on day 10-13 to human γδT cells. Murine γδT cells were activated with cytokines overnight and human γδT cells were activated with cytokines for 4 hours.

### Flow cytometry analysis

After cytokine stimulation, cells were incubated with fluorochrome-conjugated antibodies for 30 minutes at 4°C. Cells were then washed twice with staining buffer and re-suspended in staining buffer, ready to be acquired by BD LSRFortessa™ X-20 (BD Biosciences, San Jose, CA). For intracellular staining, cells were incubated with 10µg/ml Brefeldin A (Biolegend, San Diego, CA) for 4 hours. After 4 hours of incubation, cells were spun and subsequently stained for surface markers. Thereafter, cells were fixed and permeabilized with Intracellular Fixation and Permeabilization Buffer Set (eBioscience, San Diego, CA) and stained with intracellular antibodies according to manufacturer’s protocol. Analysis of FACS data was conducted with FlowJo™ software (Tree Star, Ashland, OR). The following fluorochrome-conjugated antibodies were purchased from Biolegend (San Diego, CA): PE/Dazzle™ 594 anti-mouse CD3 (17A2), PE/Cy7 anti-mouse TCR γ/δ (GL3), APC anti-mouse TCR Vγ1.1 (2.11), FITC anti-mouse TCR Vγ4 (UC3-10A6), Pacific Blue™ anti-mouse CD8a (53-6.7), PE anti-mouse NKG2D (CX5), Alexa Fluor® 700 anti-mouse IFN-γ (XMG1.2), PE/Cy7 anti-mouse TNF-α (MP6-XT22), PerCP/Cy5.5 anti-mouse TNF-α (MP6-XT22), Pacific Blue™ anti-human/mouse Granzyme B (GB11), APC anti-mouse IL-17A (TC11-18H10.1), FITC anti-mouse CD45.1 (A20), PE anti-mouse CD69 (H1.2F3), Alexa Fluor® 700 anti-mouse NK1.1 (PK136), Brilliant Violet 605™ anti-mouse Ki-67 (16A8), FITC anti-mouse CD69 (H1.2F3), Alexa Fluor® anti-mouse CD62L (MEL-14), PE anti-mouse CD44 (IM7), PE/Cy7 anti-human TCR γ/δ (B1), Brilliant Violet 605™ anti-human CD16 (3G8), PE anti-human CD25 (M-A251), BV711 anti-human NKG2D (1D11), PE/Dazzle™ 594 anti-human Perforin (dG9), Brilliant Violet 605™ anti-human IL-17A (BL168), APC-Cy7 anti-human TNF-α (Mab11), PerCP/Cy5.5 anti-human IFN-γ (B27), Pacific Blue™ anti-human TCR Vδ2 (B6), APC-Cy7 anti-human CD8 (HIT8a), PE-Dazzle™ 594 anti-human CD56 (5.1H11), Alexa Fluor 700 anti-mouse CD45 (I3/2.3), Brilliant Violet 785 anti-human IFN-γ (4S.B3), Brilliant Violet 650 anti-human TNF-α (MAb11). FITC/PE anti-human REAfinity™ TCR Vδ1 (REA173) and APC anti-human REAfinity™ TCR Vδ2 (REA771) were purchased from Miltenyi Biotec (Bergisch Gladbach, Germany); PE/Cy7 anti-mouse FasL (MFL3), PE anti-mouse Perforin (eBioOMAK-D) and PE anti-mouse IL-17A (TC11-18H10.1) were purchased from eBioscience (San Diego, CA); BUV395 anti-mouse CD45.2 (104), APC-H7 anti-mouse CD4 (GK1.5), Alexa Fluor® 647 anti-mouse IFN-γ (XMG1.2), BUV395 anti-human CD45 (HI30), BUV496 anti-human CD3 (UCHT1), BUV805 anti-human CD4 (M-T477) were purchased from BD Bioscience (San Diego, CA). Either LIVE/DEAD Fixable Yellow Dead Cell Stain Kit (Invitrogen, Waltham, MA) or DAPI (Invitrogen, Waltham, MA) was used to exclude dead cells.

### Cytotoxicity assay

The DELFIA EuTDA cytotoxicity assay (Perkin Elmer, Waltham, MA) was conducted according to manufacturer’s protocol. Target cells were labelled with fluorescence enhancing ligand, BATDA. After three washes, 5-10×10^3^ target cells were separately seeded in V-bottom plates. Following that, cytokine pre-activated γδT cells or control γδT cells were seeded in different effector to target ratios. Controls for spontaneous release, maximum release by target cells and culture medium background were also included. The effector and target cells were co-cultured for 2 hours at 37°C. The supernatant which contains the TDA released from target cells will chelate with Europium and fluoresce. The fluorescence was quantified with VICTOR^3^ plate reader (Perkin Elmer, Waltham, MA). The cytotoxicity was calculated by the following formula:

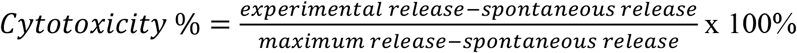

### Real-time cytotoxicity assay using xCELLigence

The xCELLigence RTCA SP (ACEA Biosciences, San Diego, CA) was used to monitor the impedance of the adherent target cells. Briefly, 50ul of RPMI-1640 was first added to each well of the 96-well E-plates (Agilent, Santa Clara, CA) and was measured as background impedance. Adherent target cells were seeded at a density of 20,000 (Hepa1-6) or 5,000 (SMMC-7721) cells/100ul to the E-plate. The plate was left in the hood at ambient temperature for 30 minutes to ensure that the cells were settled and distributed equally to the bottom of the wells before putting into a 37°C incubator. Recording of data took place every 15 minutes overnight for the target cells to attach and proliferate. Before taking out the plate for the addition of effector cells, data acquisition was paused. Pre-activated γδT cells were added at different effector to target ratios in 50µl volume. Target only controls, effector only controls and full lysis controls (0.25% Triton X) were set up. Cell index is the output of the measurement of impedance. The data were analysed with RTCA software Pro (ACEA Biosciences, San Diego, CA) which enables the determination of percentage cytolysis, by accounting for the full lysis and effector only controls. Normalisation time was set to the time before the plate was taken out for the addition of effector cells. Effector only controls were used to plot for proliferation curves.

### CD107a-degranulation assay

Activated γδT cells were seeded in round-bottom plates with FITC conjugated anti-mouse CD107a (1D4B) antibody or anti-human CD107a (H4A3) (Biolegend, San Diego, CA). γδT cells were co-cultured with A20 or K562 at a ratio of 1:1. After 1 hour of incubation, 10ug/ml Brefeldin A and 1uM monensin (Sigma-Aldrich, St. Louis, MO) were added and incubated overnight at 37°C. The next day, cells were stained with surface markers and analysed by flow cytometry.

### Murine subcutaneous melanoma model

7-9 weeks old female C57BL/6J mice were used for the subcutaneous melanoma model. 5×10^5^ B16-F10 cells in 200ul PBS were subcutaneously injected into the posterior right flank of mouse. After 10 days, when the tumor was visible to the naked eyes, it was measured every two days by calliper. The tumor volume was calculated with this formular: tumor volume (mm^3^) = (L*L*W)/2. On day 13 and day 17 post tumor implantation, 10×10^6^ control (IL-2 alone), IL-12/18, IL-12/15/18, or IL-12/18/21 pre-activated murine γδT cells generated from CD45.1-TCR-β knockout C57BL/6J mice were intratumorally injected into tumor-bearing mice. On day 21, these tumor-bearing mice were sacrificed, and spleen, tumor-draining lymph node (TDLN) and tumor were collected for lymphocyte isolation and FACS analysis.

### Murine HCC model

Murine HCC model was established with 7-9-weeks old C57BL/6J mice via hydrodynamic delivery of 1×10^6^ Hepa1-6 cells through the tail vein (1×10^6^ cells in 2ml PBS per mouse). After one week, 10×10^6^ γδT cells (control and cytokine pre-activated) (n=4-7 per group) were adoptively transferred intravenously through the tail vein once per week for two weeks. γδT cells were washed three times with PBS before adoptive transfer to remove any remaining cytokines. In the third week, mice were sacrificed and spleen and liver were harvested. Liver was weighed and tumor nodules that were formed on the surface of the liver were counted. Spleen and liver were homogenised and single cell suspensions were obtained. Specifically, lymphocytes from the liver were isolated via Percoll density centrifugation. For intracellular staining, cells were stimulated with cell activation cocktail (Biolegend, San Diego, CA) and Brefeldin A for 4 hours prior to fixing and permeabilization. Cells were then stained with flow antibodies and acquired by BD LSRFortessa™ X-20 (BD Biosciences, San Jose, CA).

For combination treatment of anti-PD-L1 antibody and adoptive γδT cell transfer, one week after the establishment of the tumor model, mice received either IL-12/18/21 pre-activated CD45.1 γδT cells only, anti-PD-L1 antibody only or both. γδT cells were transferred on day 7 and 14. 100µg *InVivo*MAb rat IgG2b isotype control (LTF-2, Bio X Cell, Lebanon, NH) or *InVivo*MAb anti-mouse PD-L1 (B7-H1) (10F.9G2, Bio X Cell, Lebanon, NH) were injected on day 7, 14 and 18. Mice were sacrificed at the end of the third week for flow analysis.

### CD8^+^T cell depletion in murine HCC model

Murine HCC model was established as described above. For CD8^+^T cell depletion, mice were intraperitoneally injected *InVivo*MAb anti-mouse CD8 antibody (2.43, 300ug/mouse, Bio X Cell, Lebanon, NH) on day 1, day 8 and day 15. *InVivo*MAb rat IgG2b (LTF-2, 300ug/mouse, Bio X Cell, Lebanon, NH) was injected as isotype control.

### Humanized mouse model

NOD-SCID IL2Rγ^-/-^ (NIKO) mice were generated and bred in the animal facility at A*STAR, Biological Resource Centre (BRC). Within 72 hours of birth, neonate mice were sub-lethally irradiated (100 rads) and injected extrahepatically with 1×10^5^ human CD34^+^ hematopoietic stem cells (HSCs). At 12-weeks post-transplantation, human immune cell reconstitution levels in the peripheral blood of mice were determined via flow cytometry. The percentage of human CD45^+^ cells was cut off at 55% out of the sum of human CD45^+^ and mouse CD45^+^ cells(42). Humanized murine HCC model was established through subcutaneous engraftment of 1×10^6^ HepG2 cells resuspended in 100ul of PBS. After 1 week, control (IL-2 alone) or IL-12/18/21 pre-activated human γδT cells expanded by anti-TCR pan-γδ clone IMMU510 antibody or Zol were administered via intravenous injection at indicated times (Fig. 5A). To overcome the IL-12/18-induced apoptosis, human γδT cells were pre-treated with p-JNK inhibitor (SP600125: 20uM, Invivogen, San Diego, CA) for 30min followed by IL-12/18/21 stimulation for 4 hours(43). Tumor size was measured by calliper every 3 days. The tumor volume was calculated with this formular: tumor volume (mm^3^) = (L*L*W)/2. On day 28, the tumor-bearing humanized mice were sacrificed, and the tumors were harvested for weight measurement, lymphocyte isolation and FACS analysis. Mice were bred and housed under specific pathogen free conditions. All animal experimental procedures were conducted in accordance with the guidelines of Agri-Food and Veterinary Authority and the National Advisory Committee for Laboratory Animal Research of Singapore and were approved by the International Animal Care and Use Committee (IACUC), A*STAR (BRC #181367). Written informed consent was obtained from donors. This study was conducted in accordance with the approval of A*STAR-IRB (#2020-040).

### Co-culture of CD8^+^T cells and γδT cells

Murine γδT cells were pre-activated with cytokines as previously described. γδT cells were washed three times before co-culturing at a ratio of 1:5 with CD8^+^T cells isolated from tumor-bearing mice using CD8a (Ly-2) microbeads through magnetic activated cell sorting (MACS) (Miltenyi Biotec, Bergisch Gladbach, Germany). Tumor cell lysate (TCL) derived from Hepa1-6 cells were added to the co-culture system at a ratio of 1 γδT cell to 3 tumor cell lysates in 96-well round-bottom plate. Transwell plate (0.4um pore size polycarbonate membrane, Sigma-Aldrich, St. Louis, MO) was set up to block cell-cell contact between CD8^+^T cells and γδT cells. CD8^+^T cells were seeded in the bottom compartment and γδT cells and tumor cell lysate were added in the upper compartment. After 3 days of co-culture in 37°C incubator, surface markers and cytokine production by CD8^+^T cells were measured and analysed by flow cytometry.

### In vivo proliferation assay

Donor γδT cells from CD45.1 TCRβ knockout mice were pre-activated with cytokines overnight. Cells were washed and labelled with 25µM CellTrace™ CFSE Cell Proliferation kit (Invitrogen, Carlsbad, CA) at a concentration of 50 × 10^6^ cells/ml. CFSE-labelled cytokine pre-activated and control γδT cells were transferred into Hepa1-6 tumor-bearing mice. After four days, mice spleen and liver were harvested, stained, and analyzed with FACS. Individual CFSE generations of the transferred γδT cells could be visualized and the proliferation index could be calculated using FlowJo software (Tree Star, Ashland, OR).

### Apoptosis of γδT cells

Cells were stained with FITC conjugated Annexin V (Biolegend, San Diego, CA) and DAPI and analysed with flow cytometry. Apoptotic cells are defined as Annexin V^+^/DAPI^-^.

### Statistical analysis

All statistical analyses were performed with GraphPad Prism version 7.0 (GraphPad Software, San Diego, CA). Multiple comparisons were performed using ordinary one-way ANOVA with Dunnett’s post-hoc test when comparing a control or two-way ANOVA with Tukey’s post-hoc test when comparing among all groups. Data were shown as mean ± SEM, and *p* <0 .05 was considered statistically significant. * indicates *p* < 0.05, ** indicates *p* < 0.01, *** indicates *p* < 0.001. Non-significant differences were not annotated.

## Results

### Cytokine pre-activation promoted murine γδT cell proliferation, perforin expression and IFN-γ production

To investigate which cytokine combination could promote murine γδT cell activation and cytokine production, we generated murine γδT cells in the presence of single or different combinations of cytokines for 16 hours *in vitro*. The expressions of activation markers and cytokine productions were measured. γδT cells treated with IL-18 and the combinations which consist of IL-18 led to an increase in CD69 expression (Fig. 1A). CD25 was upregulated by IL-12 (Fig. 1B). IL-12 and IL-18 synergistically elicited a significant increase in IFN-γ production as compared to single cytokine treatment (Fig. 1C). Thus, IL-12 and IL-18 are important cytokines that could potentially enhance the anti-tumor function of γδT cells. IL-15 and IL-21 were kept in the combination because it is reported that IL-15 could increase the proliferation and cytotoxic capacity of human γδT cells(34), while IL-21 is able to enhance survival and anti-tumor activity of CD8^+^ T cells(44). Subsequent experiments mainly focused on the four combinations of cytokines: IL-12/18, IL-12/15/18, IL-12/18/21 and IL-12/15/18/21. Murine γδT cells were monitored for 48 hours real-time upon cytokine treatment. All cytokine combinations greatly promoted γδT cell proliferation compared to control group (Fig. 1D). The proportions of the major γδT subsets found in mouse, Vγ1 and Vγ4, were not affected in response to the cytokine treatment (Fig. 1E). Vγ1^+^ cells were approximately 40% and Vγ4^+^ cells were 30% of the total γδT cell population. All cytokine combinations induced a slight increase in the percentages of CD25 expression compared to control (Fig. 1F). FasL was significantly upregulated in all combinations (Fig. S1A). Prominently, cytokine combinations consisting of IL-21 resulted in a lower FasL expression. Cytokines did not greatly affect the expression of NKG2D expression with a slight increase with IL-12/18/21 and IL-12/15/18/21 treatments (Fig. S1B). NKG2D mRNA expression was not affected by the cytokine treatment either (Fig. S1D). Since γδT cells can express NK receptors to engage tumor recognition and lysis, we also examined the expressions of other NK activating receptors (NKG2C, KLRA4 and CRTAM) and cytotoxic receptors (CD226, CD244, CD16 and NCR1) on murine γδT cells upon cytokine stimulation (Fig. S1D). The results showed that only CRTAM was significantly upregulated by IL-12/18/21. NKG2C, KLRA4, CD226, CD16, CD244 as well as NCR1 were significantly downregulated by IL-12/18/21 treatment. All cytokine combinations resulted in higher perforin levels as compared to control and the inclusion of IL-21 in the combination further boosted it (Fig. 1G). Granzyme B was constitutively expressed at a high level in all groups (Fig. 1H). IL-17A production was reduced when γδT cells were treated with cytokine combinations while TNF-α levels were increased (Fig. S1C and 1I). As expected, all cytokine combinations dramatically increased IFN-γ production in both percentages and expression levels (Fig. 1J). Thus, cytokine combinations of IL-12/18, IL-12/15/18, IL-12/18/21 and IL-12/15/18/21 promoted γδT cell proliferation, perforin expression and IFN-γ production.

**Figure 1:**
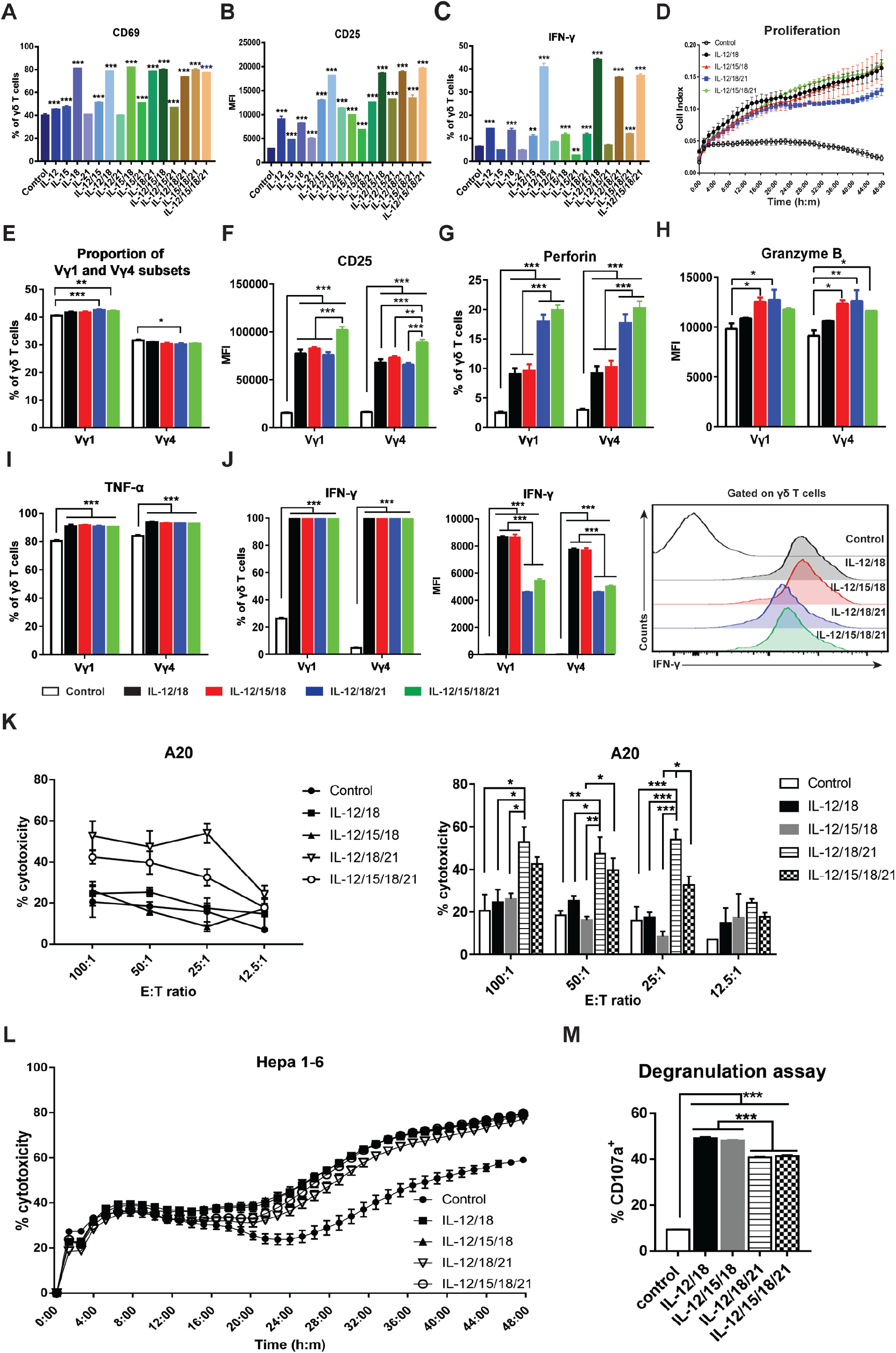
Cytokine pre-activation promoted murine γδT cell proliferation, perforin expression, IFN-γ production, and cytotoxicity *in vitro*. (A-C) Flow cytometry analysis of the surface expressions and cytokine productions by murine γδT cells to screen the effect of individual cytokines and their combinations (IL-12:10ng/ml, IL-15:100ng/ml, IL-18:50ng/ml, IL-21:30ng/ml). The percentages or MFI of the activation markers CD69 (A) and CD25 (B). (C) The percentage of IFN-γ-producing γδT cells. (D) Growth curves of cytokine pre-activated γδT cells *in vitro*. 50,000 γδT cells per well were seeded in 96-well E-plate and the cell index was monitored for 48 hours using xCELLigence. (E) The proportions of Vγ1 and Vγ4 subsets upon cytokine treatment. (F-J) The expressions of activation markers and cytokine productions of γδT cells upon cytokine treatment. Murine γδT cells (1×10^6^/ml) were stimulated with the four combination of cytokines (white: Control-IL-2 only, black: IL-12/18, red: IL-12/15/18, blue: IL-12/18/21, green: IL-12/15/18/21) for 16 h, then the expressions of CD25 (F), Perforin (G), Granzyme B (H), TNF-α (I) and IFN-γ (J) in Vγ1 and Vγ4 subsets were measured by flow cytometry. (K) Cytotoxicity assay was performed by co-culturing murine γδT cells that were pre-activated with respective cytokine combinations for 16h with BATDA-labelled target cells A20 at effector to target (E:T) ratio ranging from 100:1 to 6.25:1 for 2h. Killing capacity of the cytokine pre-activated murine γδT cells were measured by the amount of BATDA released by target cells when lysed. Right panels indicate statistical significance of cytotoxicity compared among groups. (L) Real-time cytotoxicity of cytokine pre-activated γδT cells against Hepa1-6 cells was monitored for 48 hours with E:T ratio of 2.5:1 using xCELLigence. (M) The expression of CD107a by cytokine pre-activated γδT cells when co-cultured with A20. Results shown (A-M) are representative of at least 3 independent experiments. The graph displays mean ± SEM. Significance was determined by one-way ANOVA with Dunnett’s multiple comparison test (A-C and M) and two-way ANOVA with Tukey’s multiple comparison test (D-L). *** *p* <0.001 ** *p* <0.01 * *p* <0.05.

### Cytokine pre-activated murine γδT cells exhibited enhanced cytotoxicity against tumor cells

To determine whether the cytokine pre-activation could promote the cytotoxicity of murine γδT cells against tumor cells, we first examined short-term cytotoxicity against tumor targets of A20 (murine B-cell lymphoma cell line) and Hepa1-6 (murine hepatoma cell line) cells. Murine γδT cells treated with two cytokine combinations, namely IL-12/18/21 and IL-12/15/18/21, significantly increased their cytotoxicity against A20 compared to control γδT cells (Fig. 1K). All cytokine combinations contributed to an increased cytotoxicity against Hepa1-6 targets, with IL-12/18/21 and IL-12/15/18/21 combinations being the most effective (Fig. S1E). Adopting a label-free electrical impedance-based assay which is capable of monitoring the killing activity of γδT cells in real-time and for longer durations, we explored the killing kinetics of cytokine-activated γδT cells. Overall, all cytokine combinations promoted killing capacity of γδT cells as compared to the control (Fig. 1L). To extend these observations, we analysed the amount of secretory lysosomes by measuring the expression of CD107a, which correlates with cytotoxicity(45). Cytokine-activated murine γδT cells had higher CD107a expression compared to control γδT cells in the presence of A20 (Fig. 1M). Altogether, these results demonstrated that cytokine pre-activated murine γδT cells have enhanced killing capacity and may exert more effective anti-tumor activity upon adoptive transfer.

### Cytokine pre-activation promoted CD25 expression and cytokine production in human γδT cells

To determine whether cytokine pre-activation has similar effects on human γδT cells, we examined the effects of cytokine combinations of IL-12/18, IL-12/15/18, IL-12/18/21 and IL-12/15/18/21 on human γδT cells that were expanded by Zol. CD25 was highly upregulated upon cytokine activation (Fig. 2A). NKG2D and CD16 expressions were not greatly influenced by the cytokine treatment (Fig. S2A and S2B). IL-12/18/21 significantly increased the expression of CD56 on Zol-expanded human Vγ9Vδ2 T cells but not DNAM-1 (Fig. S2D). The expression level of NKp46 was very low. Perforin and granzyme B were constitutively expressed at a high level and not affected by the cytokine treatment (Fig. 2B and 2C). TNF-α and IFN-γ productions were significantly increased by all cytokine combinations (Fig. 2D and 2E).

**Figure 2:**
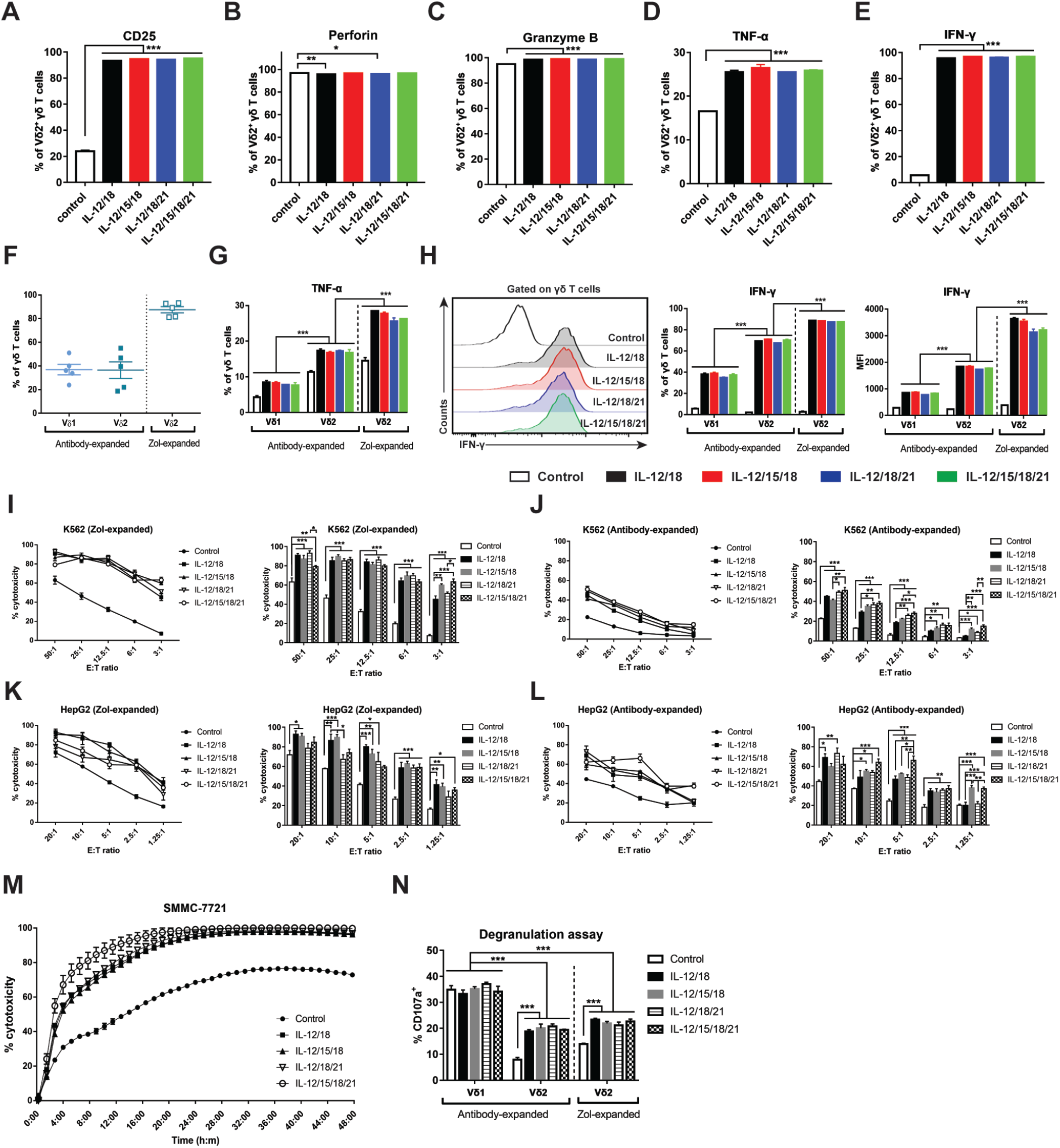
Cytokine pre-activation promoted human γδT cell CD25 expression, cytokine production and cytotoxicity against tumor cells *in vitro*. (A-E) Flow cytometry analysis of activation marker expressions and cytokines produced by Zol expanded human γδT cells at 4h upon cytokine activation (IL-12:10ng/ml, IL-15:10ng/ml, IL-18:10ng/ml, IL-21:10ng/ml). Percentages of human Vδ2^+^ γδT cells expressing CD25 (A) and production of Perforin (B), Granzyme B (C), TNF-α (D) and IFN-γ (E). Two different expansion methods, using either immobilized pan-γδTCR antibody or Zol were evaluated based on Vδ1^+^ and Vδ2^+^ subsets. (F) Percentage of Vδ1^+^ and Vδ2^+^ subsets derived from the antibody-based expansion and Zol expansion. Data is obtained from 5 donors. (G and H) The productions of TNF-α and IFN-γ by cytokine pre-activated human γδT cells. Antibody-expanded and Zol expanded (Zol-expanded) human γδT cells (1х10^6^/ml) were stimulated with cytokine combinations (white: Control-IL-2 only; black: IL-12/18; red: IL-12/15/18; blue: IL-12/18/21; green: IL-12/15/18/21) for 4 h, then the expressions of TNF-α (G) and IFN-γ (H) in Vδ1^+^ and Vδ2^+^ γδT cells were measured by flow cytometry. Zol-expanded human γδT cells from healthy donors with Vδ2^+^>80% purity were used in these experiments. (I-L) Zol-expanded and antibody-expanded human γδT cells were activated with cytokine combinations for 4h before co-culturing with BATDA-labelled target cells K562 (I-J) or HepG2 (K-L) for 2h. Killing capacity of the activated human γδT cells was assessed by the amount of BATDA released by target cells when lysed. Right panels indicate statistical significance of cytotoxicity compared among groups. (M) Real-time cytotoxicity of cytokine pre-activated Zol-expanded human γδT cells against SMMC-7721 cells was monitored for 48 hours with E:T ratio of 5:1 using xCELLigence. (N) The expression of CD107a by cytokine pre-activated human γδT cells when co-cultured with K562. Zol-expanded human γδT cells from healthy donors with Vδ2^+^>80% purity were used in these experiments. Results shown (A-H) are representative of at least 3 independent experiments. The graph displays mean ± SEM. Significance was determined by one-way ANOVA with Dunnett’s multiple comparison test when comparing between control and cytokine pre-activated human γδT (A-E) and two-way ANOVA with Tukey’s multiple comparison test (F-N). *** *p* <0.001 ** *p* <0.01 * *p* <0.05.

The expansion of γδT cells using immobilised anti-γδTCR antibody could generate polyclonal γδT cells from PBMC(46). To compare the effect of cytokine pre-activation on different subsets of human γδT cells, we expanded human γδT cells using anti-γδTCR antibody and showed that this method yielded two major subsets with high inter-individual variations among donors: Vδ1^+^ (25-50%) and Vδ2^+^ (20-55%) (Fig. 2F). On the other hand, Zol-expanded γδT cells were mostly Vδ2^+^ (80-93%). In antibody-expanded γδT cells, CD16 expression was significantly higher in Vδ2^+^ than Vδ1^+^ subset (Fig. S2C). The expressions of NKp46, DNAM-1 and CD56 were comparable between antibody-expanded Vδ2^+^ and Vδ1^+^ subsets and they were not affected by cytokine treatment (Fig. S2D). Cytokine combination treatment promoted TNF-α and IFN-γ production in all human γδT cell subsets. However, Vδ2^+^ cells had higher levels of TNF-α and IFN-γ production than Vδ1^+^ subset. When compared between the two expansion methods, Zol-expanded Vδ2^+^ cells had higher CD16, TNF-α, and IFN-γ expressions than antibody-expanded Vδ2^+^ cells (Fig. 2G, 2H and S2C). Overall, cytokine combinations promoted the activation and cytokine production of human γδT cells expanded by either Zol or anti-γδTCR antibody.

### Cytokine pre-activation enhanced the cytotoxicity of human γδT cells against tumor cells

We next examined whether cytokine pre-activation could promote the cytolytic activity of human γδT cells. We compared the cytotoxicity of Zol-expanded γδT cells and antibody-expanded γδT cells against tumor cells upon cytokine treatment. We assessed the cytotoxic effect of cytokine pre-activated γδT cells on K562, a human chronic myeloid leukemia cell line, HepG2, and SMMC-7721 which are human hepatoma cell lines (Fig. 2I-2N). All cytokine combinations promoted the killing capacity of γδT cells against K562 cells compared with the control at all E:T ratios (Fig. 2I and 2J). While antibody-expanded γδT cells exhibited similar trend as Zol-expanded γδT cells upon cytokine treatment, the cytotoxic activity of antibody-expanded γδT cells was lower compared to Zol-expanded γδT cells (Fig. 2I and 2J). In killing solid tumor targets HepG2, cytokine treatment also enhanced the cytolytic function of γδT cells with Zol-expanded γδT cells having higher cytotoxicity compared to antibody-expanded γδT cells (Fig. 2K and 2L). In real-time long-term killing assay, all cytokine combinations boosted the killing capacity of human γδT cells against SMMC-7721 cells (Fig. 2M). Cytokine-treated human γδT cells had increased CD107a expression in the presence of K562 for the Vδ2^+^ subsets but not for Vδ1^+^ subset, which had an initial higher level of CD107a expression (Fig. 2N). Collectively, these data demonstrated that like murine γδT cells, cytokine pre-activation enhanced the cytotoxicity of human γδT cells against tumor cells.

### IL-12/18/21 pre-activated γδT cells inhibited tumor growth in murine melanoma and HCC models

To examine the *in vivo* efficacy of adoptively transferred cytokine pre-activated γδT cells, mice were challenged subcutaneously with B16-F10 cells followed by γδT cells adoptive transfer on day 13 and day 17 post tumor implantation (Fig. 3A). Since the cytokine combination of IL-12/15/18/21 were not superior in upregulation of cytokine production or killing capacity of γδT cells compared with the other three combinations examined, we decided to eliminate this group in the *in vivo* experiments. The intratumoral injection of IL-12/15/18 or IL-12/18/21 pre-activated CD45.1^+^ γδT cells inhibited tumor growth compared to PBS control or the control γδT cells (Fig. 3B). While IL-12/15/18 or IL-12/18/21 pre-activated murine γδT cells only significantly reduced tumor volume at the end time point compared to PBS control (Fig. 3C), their tumor weights were significantly reduced compared to those of either PBS or control γδT cells (Fig. 3D). By tracking the distribution of adoptively transferred γδT cells via CD45.1 staining, we found that the percentages of transferred γδT cells were significantly increased at the tumor site and spleen upon cytokine pre-activation (Fig 3E and Fig. S3A). Cytokine pre-activated γδT cells also displayed a trend of increased infiltration in the TDLN compared with control γδT cells (Fig. S3B). The CD107a expression on the cytokine pre-activated γδT cells also showed a trend of increase (Fig. S3C), indicating that these cells maintained enhanced cytotoxicity *in vivo*.

**Figure 3:**
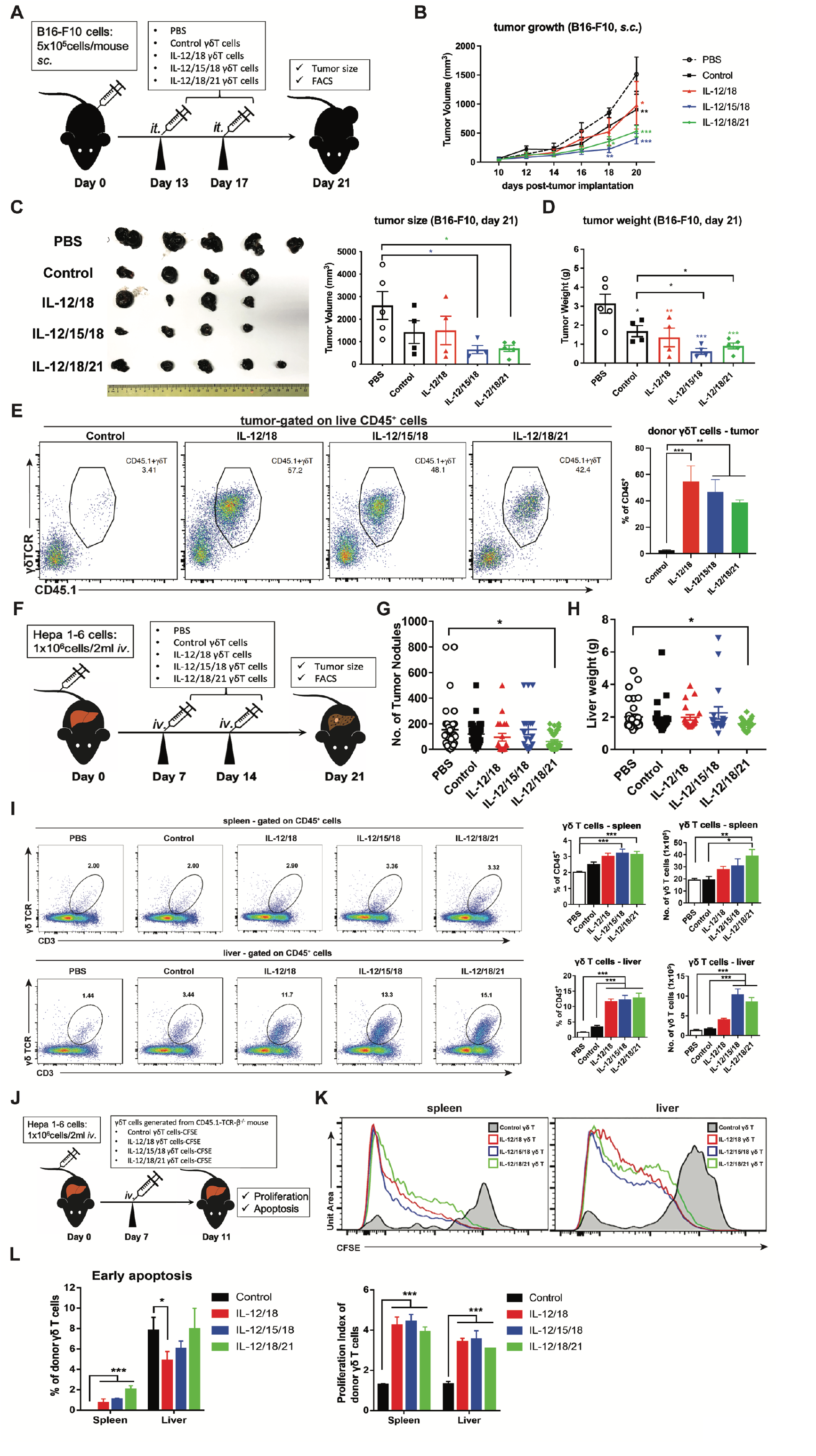
Adoptive transfer of cytokine pre-activated murine γδ T cells inhibited tumor growth in murine subcutaneous melanoma and orthotopic HCC models. (A) C57BL/6 female mice were subcutaneously injected with 5×10^5^ B16-F10 cells at the right hind limb. Tumor size was measured on alternate days from day 10. Two adoptive transfers of 10×10^6^ γδT cells that were pre-activated with three cytokine combinations, IL-12/18, IL-12/15/18, IL-12/18/21, γδT cells treated with IL-2 only or PBS (n=4-5 per group) were injected via the intratumoral route at days 13 and 17. (B) Tumor growth curve across three weeks in the groups receiving PBS: Black circle, Control: Black square, IL-12/18: red triangle, IL-12/15/18: blue triangle and IL-12/18/21: green diamond. Tumor growth was monitored every other day for a period of 20 days. (C) Images of tumor mass from each mouse and bar graph showing the endpoint tumor volumes. (D) Bar graph showing the weight of tumor mass. (E) Representative flow plots showing the CD45.1^+^ γδT population found in tumor infiltrating lymphocytes and bar graph showing the percentages of CD45.1 γδT cells in the tumor. (F) HCC mouse model was established by intravenously injecting 1×10^6^ Hepa1-6 cells with hydrodynamic cell delivery method. Two doses of control γδT cells, IL-12/18, IL-12/15/18 or IL-12/18/21 pre-activated γδT cells (10×10^6^ cells/mouse) were adoptively transferred intravenously into mice on day 7 and 14. Mice were sacrificed on day 21. (G) Tumor nodules and (H) Liver weight of mice adoptively transferred IL-12/18, IL-12/15/18, IL-12/18/21 pre-activated, or control γδT cells. PBS was injected as negative control. Data was pooled from 7 independent experiments, each group n=4-7 per experiment. (I) Representative plots showing the population of transferred γδT cells (CD3^+^ γδ TCR^+^) in the spleen (top panel) and liver (bottom panel). Right panels indicating the percentage and number of γδT cells in the spleen and liver after adoptive transfer. (J) γδT cells from CD45.1-C57BL/6 mouse were pre-activated with cytokine cocktails overnight before being labeled with CFSE and transferred into tumor-bearing mice on day 7 after hydrodynamic transfer of tumor cells. 4 days after transfer, spleen and liver were harvested and analyzed on the adoptive transferred γδ T cells’ proliferation and apoptosis.(K) Top panel: FACS profiles of transferred CFSE-labeled γδ T cells in spleen and liver (control-γδ T: black line; IL-12/18-γδ T: red line; IL-12/15/18-γδ T: blue line; IL-12/18/21-γδ T: green line). Bottom panel: bar graph showing the proliferative index of adoptively transferred γδ T cells. (L) Early apoptosis of transferred γδT cells from the spleen and liver were measured by staining Annexin V and DAPI (*n* = 3 mice per group). The data shown are representative of two experiments with 3–5 mice per group for each experiment. The graph displays mean ± SEM. Significance was determined by one-way ANOVA with Dunnett’s multiple comparison test. *** *p* <0.001 ** *p* <0.01 * *p* <0.05.

To evaluate whether the enhanced anti-tumor function of cytokine pre-activated γδT cells was tumor model specific, we established a murine orthotopic HCC model by intravenously injecting Hepa1-6 cells via hydrodynamic cell delivery as previously described(47,48). Two infusions of control γδT cells, IL-12/18, IL-12/15/18 or IL-12/18/21 pre-activated γδT cells (10^7^ cells/mouse) were performed intravenously on Day 7 and 14. The number of tumor nodules and liver weight were measured on day 21 (Fig. 3F). IL-12/18/21 pre-activated γδT cell infusion significantly reduced the number of tumor nodules and liver weight compared with PBS control (Fig. 3G and 3H). The IL-12/18 or IL-12/15/18 pre-activated γδT cell infusion slightly reduced tumor growth, but the difference was not statistically significant. In mice that received IL-12/18/21 pre-activated γδT cells, the percentages and numbers of γδT cells in spleen were significantly increased (Fig. 3I). Mice that received IL-12/15/18 and IL-12/18/21 pre-activated γδT cells were found to have higher percentages and numbers of γδT cells in the liver as compared to mice receiving control γδT cells (Fig. 3I). The adoptive transfer of cytokine-activated γδT cells did not affect the proportions of CD4^+^T, CD8^+^T cand NK cells in the liver (Fig. S3D). The results indicated that cytokine pre-activated γδT cells, especially the IL-12/15/18 and IL-12/18/21 pre-activated γδT cells, may survive longer compared to control γδT cells *in vivo*.

To assess whether the increased number of adoptively transferred γδT cells was due to the elevated proliferation of the infused cytokine-activated γδT cells, we transferred CFSE-labelled γδT cells from congenic CD45.1 mouse into tumor-bearing CD45.2 mice on day 7 after tumor establishment (Fig. 3J). CFSE expression of the transferred γδT cells was analyzed on day 11. All cytokine combinations significantly promoted the proliferation of adoptively transferred γδT cells compared to control γδT cells, both in the spleen and liver (Fig. 3K). Furthermore, the infused γδT cells were found to have comparable low apoptosis rate among all treatment groups (Fig. 3L).

We then examined the cytokine production of the transferred γδT cells in both spleen and liver from the murine HCC model (Fig. 4A and 4B). IL-12/18/21 pre-activated γδT cells exhibited significantly increased production of IFN-γ in spleen. Both IL-12/15/18 and IL-12/18/21 pre-activated γδT cells produced significantly higher levels of IFN-γ in liver (Fig. 4A and 4B). The adoptive transfer of IL-12/15/18 pre-activated γδT cells significantly increased the numbers of IFN-γ producing and TNF-α producing CD8^+^ T cells in liver (Fig. 4C and 4D) but did not affect the IFN-γ or TNF-α production of CD4^+^T and NK cells (Fig. S3E). The percentages and numbers of IFN-γ-producing and TNF-α-producing CD8^+^T cells in spleen and liver also showed an increasing trend under the treatment of IL-12/18/21 pre-activated γδT cells. Consistently, adoptively transferred IL-12/15/18 or IL-12/18/21 pre-activated γδT cells also maintained elevated production of IFN-γ in TDLN in B16-F10 subcutaneous model (Fig. S4A). However, the percentage of IFN-γ-producing γδT cells was comparable in tumor among all groups (Fig. S4B). Compared to control γδT cells, the adoptive transfer of IL-12/15/18 or IL-12/18/21 pre-activated γδT cells led to the augmented production of IFN-γ by endogenous CD8^+^T cells in spleen and TDLN but not tumor in the murine melanoma model (Fig. S4C-E). Thus, our findings indicated that cytokine pre-activation could promote the proliferation of γδT cells *in vivo*. IL-12/15/18 or IL-12/18/21 pre-activation not only upregulated IFN-γ production of the adoptively transferred γδT cells *in vivo*, but also enhanced the anti-tumor function of endogenous CD8^+^ T cells via promoting the productions of IFN-γ and TNF-α.

**Figure 4:**
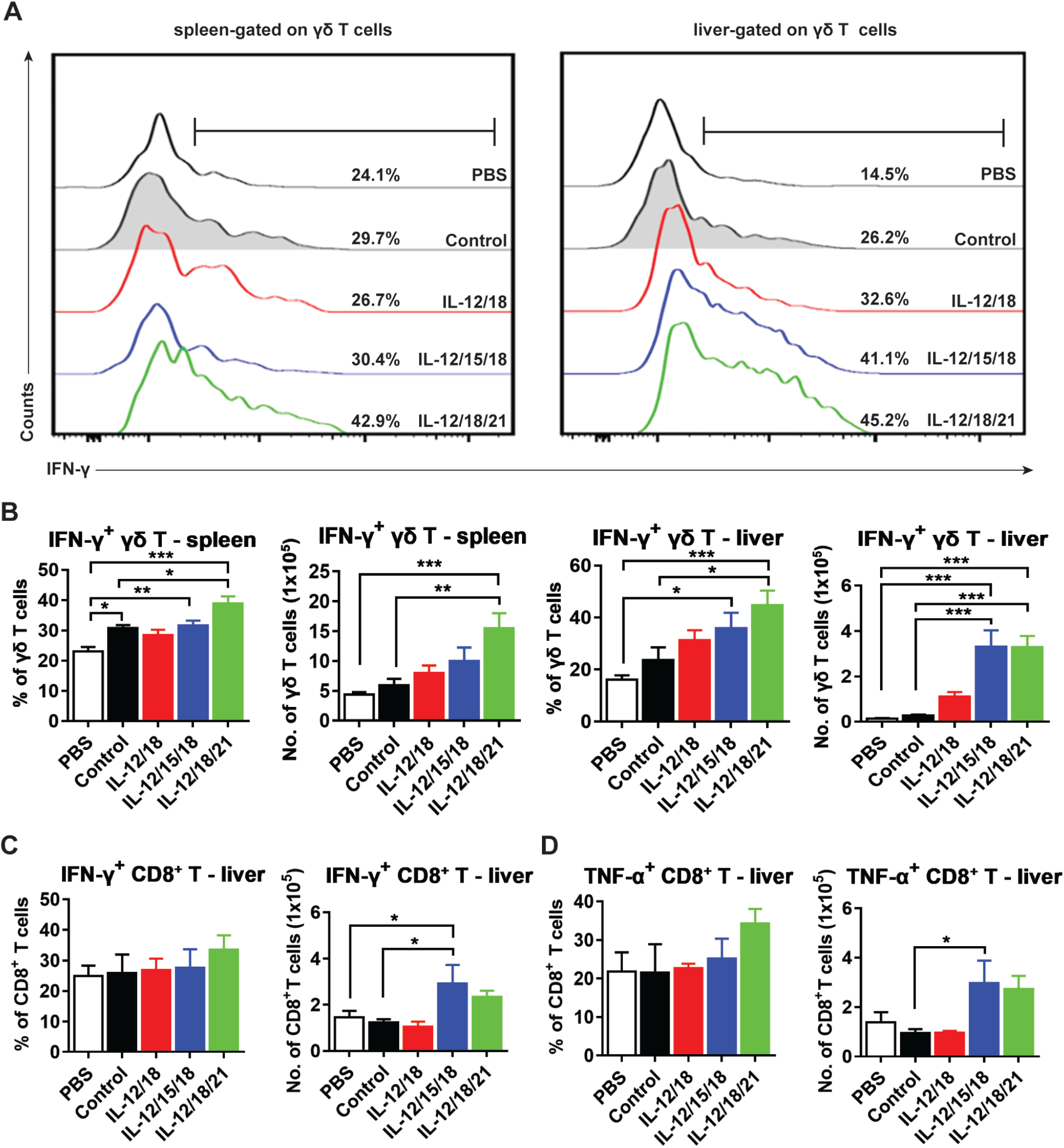
Adoptively transferred cytokine pre-activated γδT cells produced higher levels of IFN-γ and promoted cytokine production by endogenous CD8^+^ T cells *in vivo*. (A) Representative histogram plots of IFN-γ production by γδT cells gated on γδTCR^+^ in the spleen (left panel) and liver (right panel). (B) The percentages and numbers of IFN-γ-producing γδT cells in the spleen and liver after adoptive transfer. (C) The percentage and number of IFN-γ-producing CD8^+^T cells in the liver after adoptive transfer. (D) The percentage and number of TNF-α-producing CD8^+^T cells in the liver after adoptive transfer. Results shown (A-D) are representative of 7 independent experiments (n = 18-36 mice per group). The graph displays mean ± SEM. Significance was determined by one-way ANOVA with Dunnett’s multiple comparison test. *** *p* <0.001 ** *p* <0.01 * *p* <0.05.

### IL-12/18/21 pre-activated human γδT cells inhibited tumor growth in a humanized mouse model

To determine whether cytokine pre-activation could enhance the anti-tumor function of expanded human γδT cells *in vivo*, we established humanized mice by transferring CD34^+^ hematopoietic stem cells from human cord blood to irradiated NSG mice. Then HepG2 cells were subcutaneously injected to generate humanized tumor model. A recent study reported that allogeneic Zol-expanded Vγ9Vδ2 T cells exhibited promising anti-tumor function and prolonged the survival of patients with late-stage lung or liver cancer(42). However, the efficacy of antibody-expanded human γδT cells is still unclear. Here, human γδT cells were expanded with anti-TCR pan-γδ antibody (IMMU510) or Zol from an allogeneic donor. In our previous study, we found that IL-12/18 induced rapid apoptosis of human γδT cells *in vitro*. The addition of p-JNK inhibitor (SP600125:20uM) could revive human γδT cells from IL-12/18-induced apoptosis(43). Thus, after 2-weeks of expansion, human γδT cells were pre-treated with SP600125 for 30 min followed by 4 hours’ pre-activation with IL-12/18/21. The percentages of Vδ1 or Vδ2 cells were not affected by IL-12/18/21 pre-activation (Fig. S5A). There were about 80% of Vδ2^+^ cells and 20% of Vδ2^-^ cells (including ∼5% of Vδ1^+^ cells) in antibody-expanded human γδT cells. Almost all the cells were Vδ2^+^ cells for Zol-expanded human γδT cells. Same numbers of control (IL-2 alone) or IL-12/18/21 pre-activated antibody-expanded or Zol-expanded human γδT cells were adoptively transferred into tumor-bearing humanized mice on day 7, 14, 17, 21 and 24 according to the schema (Fig. 5A). PBS was used as the negative control. We found that both IL-12/18/21 pre-activated antibody-expanded and Zol-expanded γδT cells significantly inhibited tumor growth compared to PBS control, while control human γδT cells exhibited no therapeutic effect on tumor growth with even enhanced tumor development in antibody-expanded control γδT group (Fig. 5B). In consistent with the tumor growth, both IL-12/18/21 pre-activated antibody- or Zol-expanded human γδT cells showed decreased tumor volumes at the endpoint compared to negative control (Fig. 5C-E). IL-12/18/21 pre-activated antibody-expanded human γδT cells significantly reduced the tumor volume and tumor weight compared with the control antibody-expanded human γδT cells. Similar trends were seen for IL-12/18/21 pre-activated Zol-expanded human γδT cells. The percentages of tumor-infiltrating Vδ2^+^ cells from IL-12/18/21 pre-activated antibody- or Zol-expanded human γδT cell groups were significantly higher than that from the negative control group (Fig. 5F). Control Zol-expanded human γδT cells also exhibited higher Vδ2^+^ cell infiltration in tumor. On the other hand, the percentages of Vδ1^+^ cells in the tumor were comparable among all groups (Fig. S5B). This could be due to the fact that the majority of antibody- or Zol-expanded human γδT cells were Vδ2^+^ cells. Furthermore, IL-12/18/21 pre-activated antibody-expanded human γδT cells produced more granzyme B, IFN-γ and TNF-α compared with control antibody-expanded human γδT cells in the tumor (Fig. 5G-5I). IL-12/18/21 pre-activated Zol-expanded γδT cells had higher TNF-α production than control Zol-expanded human γδT cells (Fig. 5I), with no significant increase in granzyme B or IFN-γ production (Fig. 5G and 5H).

**Figure 5:**
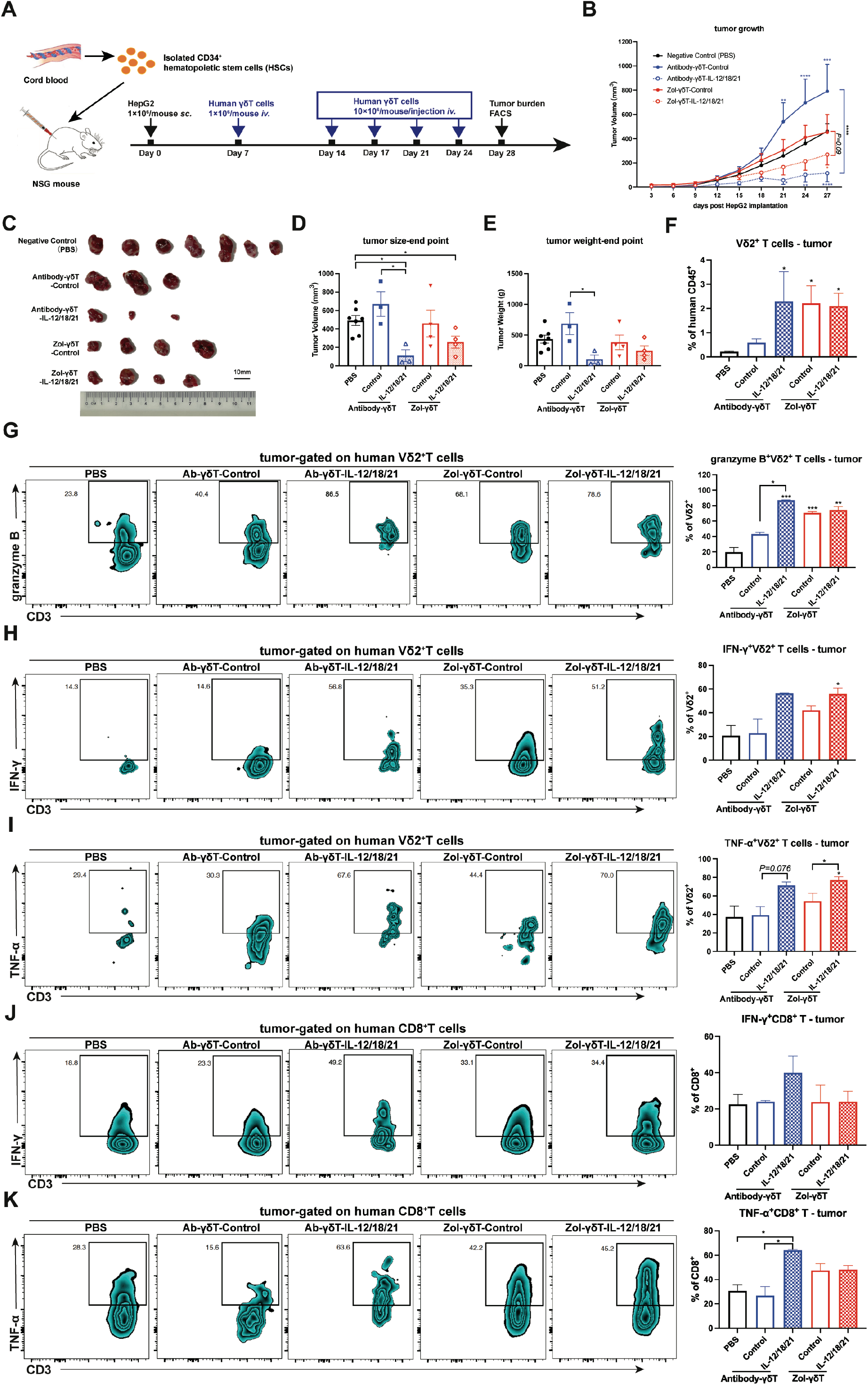
Adoptive transfer of cytokine pre-activated human γδT cells inhibited tumor growth in a humanized mouse model. (A) NOD-SCID IL2Rγ-/- (NSG) mice were sub-lethally irradiated within 72 hours of birth and injected intrahepatically with 1×10^5^ cord blood-derived human CD34^+^ hematopoietic stem cells. Mice with successful human immune cell reconstitution were subsequently used to establish the humanized murine HCC model through subcutaneous engraftment of 1×10^6^ HepG2 cells. The γδT cells were expanded with Zol or PAN-γδ TCR antibody for two weeks. γδT cells were activated for 4 hours with human recombinant IL-12 (10ng/ml), IL-18 (10ng/ml) and IL-21 (10ng/ml) before adoptive transfer. 1×10^6^ human γδT cells were administered on day 7, and subsequently 10×10^6^ human γδ T cells were administered on day 14, 17, 21 and 24 intravenously. The negative control group only received PBS intravenously. At four weeks’ post-tumor implantation, the mice were sacrificed. (B) Tumor growth curve across 4 weeks in the groups receiving PBS (Negative Control): Black solid line, Antibody-γδT-Control: Blue solid line, Antibody-γδT-IL-12/18/21: Blue dotted line, Zol-γδT-Control: Red line and Zol-γδT-IL-12/18/21: Red dotted line. Tumor were measured on alternate days. (C) Images of tumor mass from each mouse. Bar graphs showing the (D) end point tumor volume and (E) weight of tumor mass. (F) Percentages of tumor infiltrating Vδ2^+^ cells. Representative contour plots and percentages of (G) Granzyme B, (H) IFN-γ and (I) TNF-α produced by Vδ2^+^ T cells in the tumor were analyzed by flow cytometry. In the tumor, the percentage of (J) IFN-γ and (K) TNF-α produced by CD8^+^ T cells were shown. The data shown are from one experiment with 3-7 mice per group. Data shown as mean ± SEM. Significance was determined by one-way ANOVA with Dunnett’s multiple comparison test. *** *p* <0.001 ** *p* <0.01 * *p* <0.05.

To investigate whether IL-12/18/21 pre-activated human γδT cells could also promote host CD8^+^T cell functions, we examined the cytokine production of tumor-infiltrating CD8^+^T cells. The results showed that IL-12/18/21 pre-activated antibody-expanded human γδT cells significantly enhanced the TNF-α production of endogenous CD8^+^T cells in the TME with a trend of increase in IFN-γ production compared to the control antibody-expanded γδT cells (Fig. 5J and 5K). IL-12/18/21 pre-activated Zol-expanded human γδT cells did not exhibit enhancement of cytokine productions of endogenous CD8^+^T cells in the TME. These findings demonstrated that IL-12/18/21 pre-activated antibody- or Zol-expanded human γδT cells were more effective in inhibiting tumor growth than their respective control γδT cells. They had enhanced cytokine productions *in vivo*; whereas only IL-12/18/21 pre-activated antibody-expanded human γδT cells promoted the function of tumor infiltrating CD8^+^T cells in the humanized mouse model.

### IL-12/18/21 pre-activated murine γδT cells promoted CD8^+^ T cell function in a cell-cell contact dependent manner

As γδT cells are able to regulate CD8^+^ T cell function(9), which is critical for anti-tumor immunity, we investigated whether cytokine pre-activated γδT cells could directly enhance CD8^+^ T cell function. Since IL-12/18/21 combination group consistently exhibited significant reduction in tumor growth in the murine HCC models, we chose this combination in the following experiments. Murine γδT cells were pre-activated with IL-12/18/21 and co-cultured with CD8^+^T cells isolated from the spleen of tumor-bearing mice in the presence of Hepa1-6 tumor lysate. In addition, transwell inserts were used to block cell-cell contact between CD8^+^ T cells and γδT cells to determine whether the effects were dependent on cell-cell contact (Fig. 6A). IL-12/18/21 pre-activated murine γδT cells enhanced the proliferation and activation of CD8^+^T cells as shown by higher percentages of Ki67^+^ and CD69^+^ CD8^+^T cells during co-culture with direct cell contacts (Fig. 6B-C). The percentage of CD44^hi^ CD62L^-^ effector CD8^+^T cells was also increased when co-cultured with control or pre-activated γδT cells (Fig. 6D). However, when there was no cell-cell contact, the addition of γδT cells had minimal effect on the proliferation and activation of CD8^+^T cells (Fig. 6B-D). IFN-γ and TNF-α productions by CD8^+^T cells were assessed by intracellular staining. The results showed that CD8^+^T cells co-cultured with IL-12/18/21 pre-activated γδT cells produced significantly higher levels of IFN-γ and TNF-α with direct cell contact (Fig. 6E-F). However, in the absence of cell-cell contact, the effects were abrogated. Therefore, IL-12/18/21 pre-activated murine γδT cells could promote endogenous CD8^+^T cell function in a cell-cell contact dependent manner.

**Figure 6:**
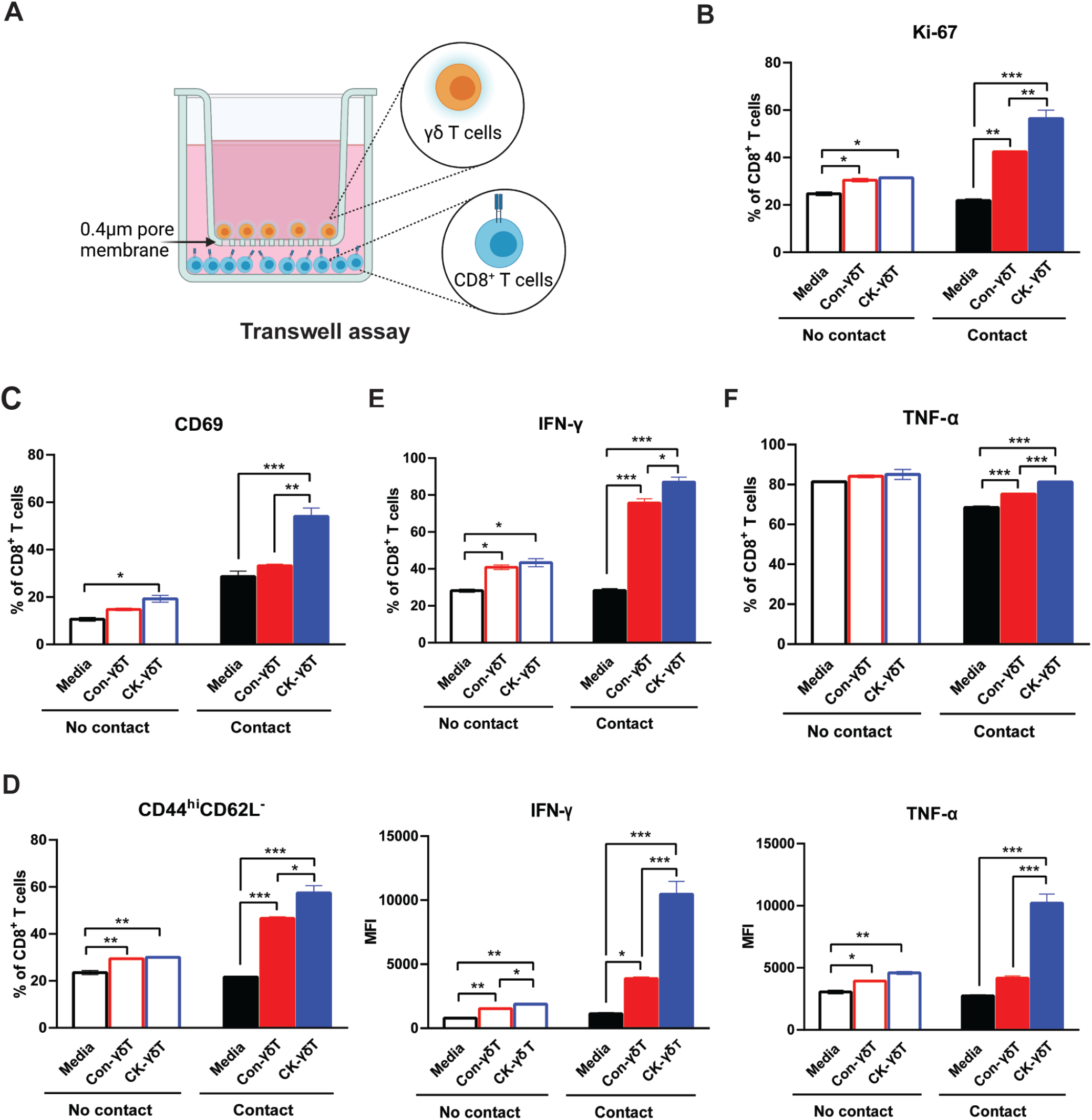
IL-12/18/21-pre-activated γδT cells promoted CD8^+^ T cell function in a cell-cell contact dependent manner. CD8^+^T cells from tumor-bearing mice were isolated and co-cultured *in vitro* with control murine γδT cells (Con-γδT) or IL-12/18/21 pre-activated murine γδT cells (CK-γδT) for three days. Hepa1-6 tumor cell lysate was added into the co-culture. (A) To prevent cell -cell contact between CD8^+^ T cells and γδT cells, a transwell assay (no-contact) was set up with CD8^+^T cells in the bottom compartment and γδT cells and tumor cell lysate were added in the upper compartment. (B) Percentage of Ki-67-expressing CD8^+^ T cells after co-culture (contact and no-contact) with no γδT, Con-γδT or CK-γδT. (C-D) Percentage of CD8^+^T cells expressing CD69 (C) and CD44^hi^ CD62L^-^ (D) effector T cells. (E-F) Percentages and MFI of IFN-γ (E) and TNF-α (F) production by CD8^+^T cells. Results shown (A-F) were representative of at least 2 independent experiments. The graph displays mean ± SEM. Significance was determined by one-way ANOVA with Dunnett’s multiple comparison test. *** *p* <0.001 ** *p* <0.01 * *p* <0.05.

### Adoptive transfer of IL-12/18/21 pre-activated γδT cells synergistically enhanced the efficacy of anti-PD-L1 antibody therapy in a CD8^+^T cell-dependent manner

Although IL-12/18/21 pre-activated γδT cell adoptive transfer significantly reduced tumor growth in murine tumor models, the efficacy was modest. PD-1 expression could hinder the anti-tumor function of γδT cells(49). Anti-PD-L1 antibody therapy has led to disease regression in many types of cancers. However, some patients developed resistance and showed limited responses. A recent study reported that sufficient IFN-γ at the tumor site can alter the gene transcription to upregulate anti-tumor function(50). The enhanced anti-tumor activity and increased IFN-γ production of cytokine pre-activated γδT cells prompted us to investigate whether adoptive transfer of IL-12/18/21 pre-activated γδT cells could overcome the resistance and have a synergistic effect with checkpoint blockade therapy. IL-12/18/21 pre-activated γδT cells were adoptively transferred on Day 7 and 14; 100µg of anti-PD-L1 per mouse were administered on Day 7, 14 and 18 (Fig. 7A). Anti-PD-L1 antibody treatment alone did not significantly influence tumor growth compared with the control group (average 4.6% decrease in tumor burden) (Fig. 7B). Again, IL-12/18/21 pre-activated γδT cell adoptive therapy alone had a significant but modest reduction in tumor development (average 41.9% reduction in tumor burden). However, the combination of the two therapeutic modalities had a drastic inhibitory effect on tumor growth (average 70.4% reduction in tumor burden). The results demonstrated a synergistic anti-tumor effect of combining IL-12/18/21 pre-activated γδT cells and anti-PD-L1 antibody therapy. Furthermore, the adoptive transfer of IL-12/18/21 pre-activated γδT cells overcame the resistance to anti-PD-L1 antibody therapy.

**Figure 7:**
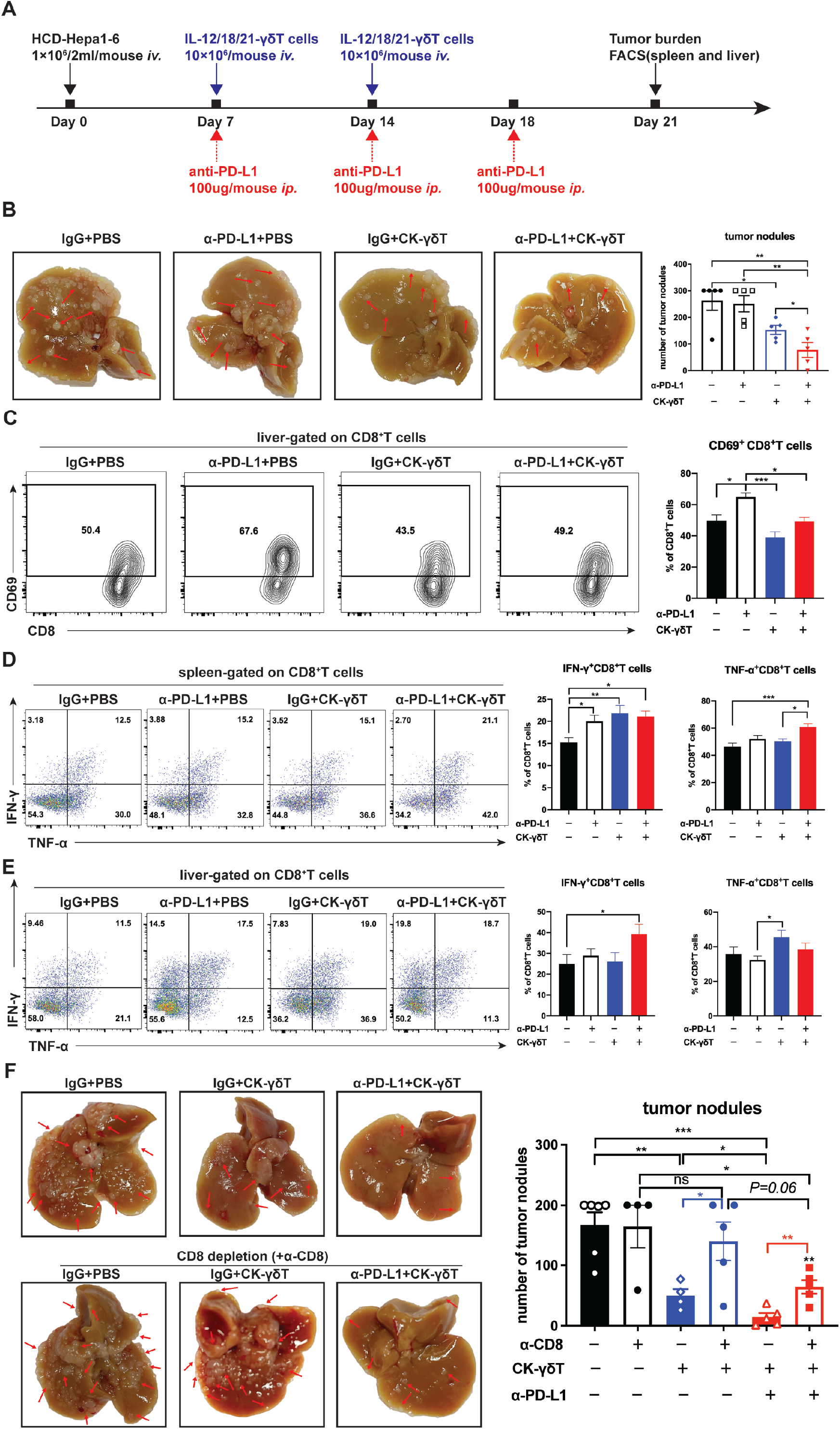
Adoptive transfer of IL-12/18/21 pre-activated γδT cells synergistically enhanced the efficacy of anti-PD-L1 antibody therapy. (A) The establishment of HCC mice model and adoptive transfer of CK-γδT (IL-12/18/21 pre-activated γδT cells) are described in Figure 3F. As a combination therapy, anti-PD-L1 antibody (100µg/mouse) was concurrently administrated intravenously on day 7, 14 and 18. Mice were sacrificed on day 21 and spleen and liver were harvested for FACS analysis. (B) Representative images of liver with tumor nodules from each group. Red arrows point to tumor nodules. (C) Representative contour plots and percentages of CD69^+^ CD8^+^T cells. (D-E) Representative FACS plots of IFN-γ and TNF-α production by CD8^+^T cells and percentages in (D) spleen and (E) liver. Results shown (A-E) are representative of 3 independent experiments (n = 5-6 mice per group). (F) *In vivo* depletion of CD8^+^T cells in C57BL/6 mice was achieved by administrating anti-CD8 antibody (300ug/mouse) via intraperitoneal route on day 1, 8 and 15 post tumor inoculation. IgG was administered as isotype control. Representative images of liver with tumor nodules from each group were shown. Red arrows point to tumor nodules. Number of tumor nodules were counted, and results were shown as bar graphs. Results shown are representative of 2 independent experiments with 4-5 mice per group. The graph displays mean ± SEM. Significance was determined by one-way ANOVA with Dunnett’s multiple comparison test. *** *p* <0.001 ** *p* <0.01 * *p* <0.05.

Anti-PD-L1 antibody therapy led to an increase in CD69^+^ expression on CD8^+^T cells in liver (Fig. 7C). CD8^+^T cells in spleen produced higher levels of IFN-γ in all treatment groups compared with the control group (Fig. 7D). The percent of TNF-α-producing CD8^+^ T cells was significantly higher in spleen of the combination treatment group and in liver of adoptive therapy group (Fig. 7D and 7E). These results suggested that the combination of anti-PD-L1 antibody therapy and IL-12/18/21-pre-activated γδT cell adoptive therapy could boost IFN-γ and TNF-α production of CD8^+^ T cells, which was critical for anti-tumor immune response.

To further explore whether the anti-tumor function of IL-12/18/21 pre-activated γδT cells or the synergistic anti-tumor effects of combination therapy was dependent on the response of endogenous CD8^+^T cells, we performed the CD8^+^T cell depletion with an anti-mouse CD8 antibody (Fig. 7F and Fig. S6). The results showed that the anti-tumor function mediated by IL-12/18/21 pre-activated γδT cells was diminished in the absence of endogenous CD8^+^T cells (Fig. 7F). The synergistic anti-tumor effect of combination therapy with checkpoint blockade was also significantly reduced when CD8^+^T cells were depleted. These results demonstrated that endogenous CD8^+^T cells were critical for the anti-tumor function of IL-12/18/21 pre-activated γδT cells either administered alone or in combination with anti-PD-L1 antibody therapy.

## Discussion

In this study, we demonstrated that cytokine pre-activated murine and human γδT cells had increased activation marker expression, cytokine production and enhanced cytotoxicity against tumor cells *in vitro*. Comparing between the two expansion methods, we found that Zol-expanded human γδT cells possessed a higher cytolytic capability than antibody-expanded human γδT cells upon cytokine pre-activation *in vitro*. The most effective cytokine combinations are IL-12/18, IL-12/15/18, IL-12/18/21, and IL-12/15/18/21. Furthermore, adoptive transfer of IL-12/18/21 pre-activated γδT cells significantly inhibited tumor growth in murine tumor models and a humanized mouse model, as well as exhibited a synergistic therapeutic effect with anti-PD-L1 antibody blockade therapy by enhancing CD8^+^T cell functions. Therefore, IL-12/18/21 pre-activation of γδT cells *ex vivo* prior to adoptive transfer provides a novel approach to boost γδT cell function *in vivo* and overcome the resistance of immune checkpoint blockade therapy in cancer patients.

Synergistic interaction between IL-12 and IL-18 on IFN-γ production by T cells and NK cells was well-demonstrated(51-53). IL-12/18 pre-activated NK cells showed enhanced anti-leukemia function without aggravating the acute graft-versus-host disease (aGVHD) in allogenic hematopoietic stem cell transplantation (allo-HSCT)(54). Our previous work also demonstrated that the adoptive transfer of murine γδT cells (especially Vγ4 cell subset) had anti-leukemia function and reduced aGVHD in a murine allo-HSCT model(55). However, the effect of different cytokine combinations on γδT cell anti-tumor function is not clear. The current study explored the possibility of expanding γδT cell *ex vivo* in the presence of cytokine combinations to improve γδT cell-based immunotherapy. While IL-12, IL-15 and IL-18 were previously used to activate NK cells for adoptive immunotherapy(40), this is the first time IL-21 was included in the cytokine combinations for the activation of γδT cells and it is interesting that IL-21 was able to downregulate IL-17A and FasL expression and upregulate perforin expression on murine γδT cells. Moreover, IL-21 seemed to enhance the cytotoxicity of γδT cells. This is in consistence with the previous study that IL-21 could upregulate lytic granules and promote cytotoxic functions and cytokine responses in human phosphoantigen-activated Vγ9Vδ2 T cells(35).

Widely used strategy to expand human Vγ9Vδ2 T cells is via stimulation with bisphosphonate or phosphoantigens such as Zol, pamidronate and BrHPP. However, expansion with phosphoantigen can only generate mainly Vδ2 subset. It was previously demonstrated that anti-γδTCR antibody could expand both Vδ1 and Vδ2 subsets(46). There were a few studies which showed higher anti-tumor efficacy using Vδ1 subset over Vδ2 subset in colon carcinoma and ovarian cancer models *in vivo*, while both subsets were shown to have equal anti-tumor activity in an *in vitro* multiple myeloma model(56). Meanwhile, some studies also reported that Vδ2 subset displayed stronger anti-tumor response in various epithelial tumors(57). In our study, cytokine pre-activation increased the expression of CD16 and productions of TNF-α and IFN-γ from both Vδ1 and Vδ2 subsets regardless of the expansion methods, while Zol-expanded γδT cells exhibited elevated cytotoxicity against tumor cells compared to antibody-expanded γδT cells *in vitro*. DNAM1 expressed by human γδT cells was reported to contribute to HCC cell lysis(58). In our study, both antibody- and Zol-expanded human γδT cells expressed high level of DNAM1. However, the IL-12/18/21 treatment did not affect the expression of DNAM1 on human γδT cells (Fig. S2D) and it even downregulated DNAM1 (CD226) mRNA expression in murine γδT cells (Fig. S1D), suggesting that DNAM1 may not play a role in the enhanced cytotoxicity of cytokine pre-activated γδT cells. For the first time, we compared the anti-tumor function of antibody- and Zol-expanded human γδT cells *in vivo*. In tumor-bearing humanized mice, neither control antibody-expanded nor Zol-expanded human γδT cells exhibited any anti-tumor effect. Interestingly, both IL-12/18/21 pre-activated antibody- and Zol-expanded human γδT cells were able to inhibit tumor growth in the humanized mouse model. These results suggest that the cytokine pre-activation strategy may be applicable to various expansion methods that are used to generate γδT cells for adoptive transfer in clinical trials. We also observed increased Vδ2^+^ γδT cells at the tumor site (Fig. 5F), suggesting that the enhanced anti-tumor effect via cytokine pre-activation was very likely to be mediated by Vδ2^+^ γδT cells.

Previous study has shown that upon microbial activation, human γδT cells could migrate to lymph nodes and transform into professional antigen presenting cells (APCs), termed γδT-APCs, which in turn induced CD4^+^T cell responses(9). Human γδT-APCs can also efficiently cross-present soluble proteins to CD8^+^T cells(59). We also found IL-12/18/21 pre-activated murine γδT cells pulsed with tumor antigens enhanced the cytokine production of CD8^+^T cells in a cell-cell contact dependent manner *in vitro* (Fig. 6). In addition, the adoptive transfer of IL-12/18/21 pre-activated murine γδT cells and antibody-expanded human γδT cells promoted the productions of IFN-γ and TNF-α of endogenous CD8^+^T cells in murine tumor models and humanized mouse model respectively (Fig. S4C, S4D, 4C, 4D, 5J and 5K). The enhanced anti-tumor function of IL-12/18/21 pre-activated γδT cells was diminished when CD8^+^T cells were depleted in murine HCC model indicating that endogenous CD8^+^T cells were critical for the anti-tumor efficacy of IL-12/18/21 pre-activated γδT cells. IL-12/18/21 pre-activated γδT cells might promote CD8^+^T cell function via their enhanced APC functions. Still, the molecular mechanism regulating the interaction between CD8^+^T cells and γδT cells needs further investigation.

Two strategies are being adopted to enhance the anti-tumor function of immune cells: boost the anti-tumor response of immune cells and remove the immunosuppression induced by the tumor microenvironment. Adoptive cell transfer and immune checkpoint blockade therapy are two representative therapies to increase the anti-tumor response of the immune system(60). Immune checkpoint blockade therapy targets immunosuppression and has had some success in various types of cancers(61). Unfortunately, only a minority of patients respond to the current immune checkpoint blockade therapy(62). There is an urgent need to develop strategies that could expand the patient cohort that can respond and increase durable responses. Combination therapy is a promising strategy to boost the outcome of immune checkpoint therapy. Adoptive cell therapy can enhance cell infiltration in the tumor microenvironment to provide sufficient immune cells when combined with immune checkpoint blockade antibody. In our study, cytokine pre-activated γδT cells showed significantly enhanced proliferation *in vitro* and *in vivo*, increasing the number of γδT cells in the TME. Furthermore, cytokine pre-activated γδT cells secreted large amounts of IFN-γ. A recent study demonstrated that IFN-γ triggered melanoma cells to increase expression of IFN pathway genes, antigen presenting machinery and T cell-attracting chemokines that drove the enhanced response to immune checkpoint blockade therapy in melanoma(50). Here, the combination of IL-12/18/21 pre-activated γδT cell adoptive transfer and anti-PD-L1 antibody therapies significantly inhibited tumor growth possibly via IFN-γ production and enhanced anti-tumor response of CD8^+^T cells in the TME. Thus, our results indicated that IL-12/18/21 pre-activation of γδT cells could be a novel strategy to enhance the anti-tumor function of adoptively transferred γδT cells and overcome the resistance to immune checkpoint blockade therapy.

## Acknowledgements

We thank Mr. Teo Guo Hui and Dr. Paul Edward Hutchinson for their assistance in flow cytometry.

## Data Availability

The authors declare that the data supporting the findings of this study are available within the article and provided as Source Data file.

## Author contributions

HY.T, Y.S, and H.L designed the research. HY.T, Y.S and Y.L performed the experiments and analyzed the data. KSM.Y and Q.C performed the humanized mice experiments. Y.M, Z.B.H and Y.Z assisted in the experiments. N.R.J.G and H.L supervised the study. HY.T, Y.S, and H.L wrote the manuscript.

