## Supplementary data for "IL-12/18/21 pre-activation enhances the anti-tumor efficacy of expanded γδT cells and overcomes resistance to anti-PD-L1 treatment"

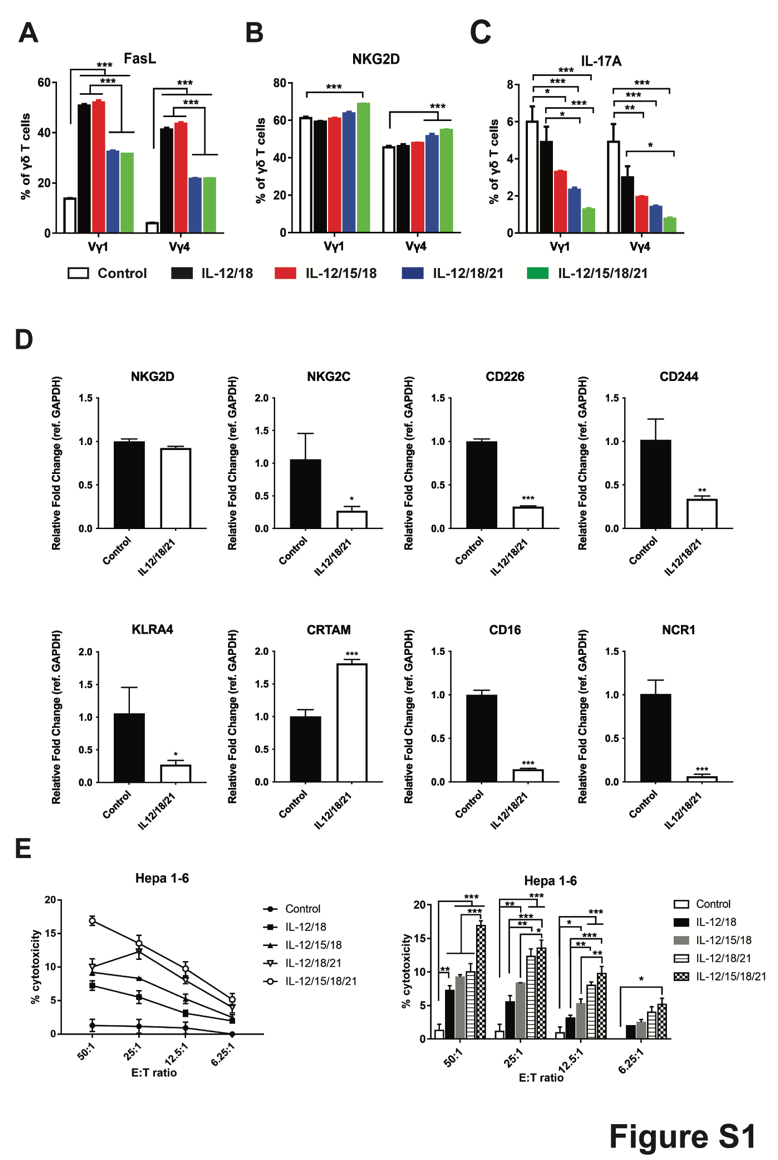

**Figure S1: Expression of NK activating receptors and in vitro cytotoxicity to Hepa1-6 cells of cytokine-preactivated mouse γδ T cells.** (A-C) The expressions of activation markers and cytokine productions of γδT cells upon cytokine treatment. Murine γδT cells (1×10^6^/ml) were stimulated with the four combination of cytokines (white: Control-IL-2 only, black: IL-12/18, red: IL-12/15/18, blue: IL-12/18/21, green: IL-12/15/18/21) for 16 h, then the expressions of FasL (A), NKG2D (B) and IL-17A (C) in Vγ1 and Vγ4 subsets were measured by flow cytometry. (D) The fold changes in the expression of NKG2D, NKG2C, CD226, CD244, KLRA4, CRTAM, CD16 and NCR1 in mouse γδ T cells after IL-12/18/21 cytokine activation compared to control were measured by real time qPCR. (E) Cytotoxicity assay was performed by co-culturing mouse γδT cells that were pre-activated with respective cytokine combinations for 16h with BATDA-labelled target cells Hepa1-6 at effector to target (E:T) ratio ranging from 50:1 to 6.25:1 for 2h. Killing capacity of the cytokine pre activated mouse γδT cells were measured by the amount of BATDA released by target cells when lysed.  Right panels indicate statistical significance of cytotoxicity compared among groups. Results shown (A-E) are representative of at least 2 independent experiments. The graph displays mean ± SEM. Significance was determined by two-way ANOVA with Tukey's multiple comparison test (A-C and E) and students’ unpaired t-test (D). *** *p* <0.001 ** *p* <0.01 * *p* <0.05.

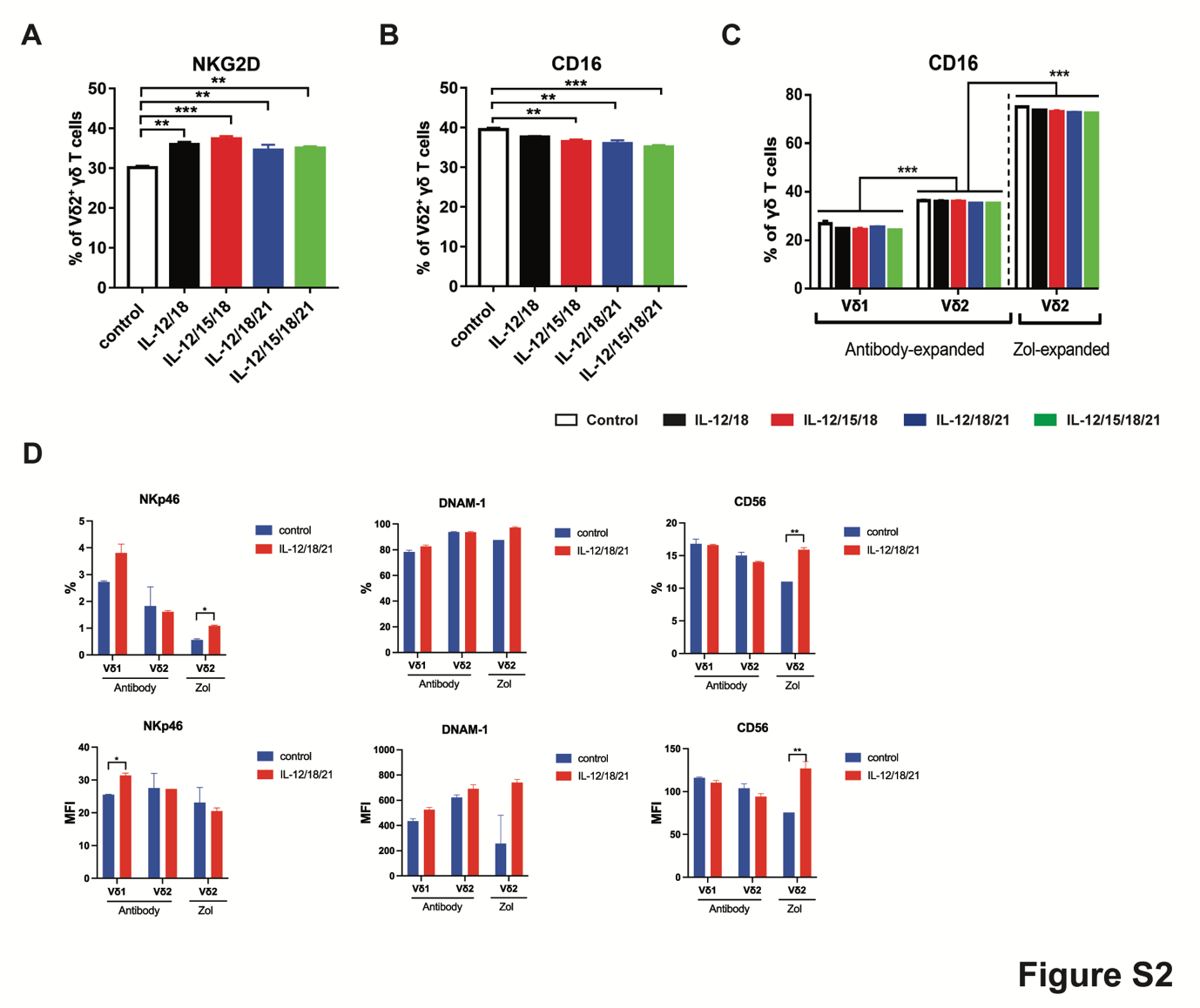

**Figure S2: Expressions of NK activating receptors in human γδ T cells.** (A and B) Percentages of Zol expanded human Vδ2^+^ γδT cells expressing NKG2D (A) and CD16 (B). Flow cytometry analysis of NKG2D and CD16 expressions by Zol expanded human γδT cells at 4h upon cytokine activation (IL-12:10ng/ml, IL-15:10ng/ml, IL-18:10ng/ml, IL-21:10ng/ml). (C) The expression of CD16 by cytokine pre-activated human γδT cells. Antibody-expanded and Zol expanded (Zol-expanded) human γδT cells (1ⅹ10^6^/ml) were stimulated with cytokine combinations (white: Control-IL-2 only; black: IL-12/18; red: IL-12/15/18; blue: IL-12/18/21; green: IL-12/15/18/21) for 4 h, then the expression of CD16 in Vδ1^+^ and Vδ2^+^ γδT cells were measured by flow cytometry. (D) The expression of NKp46, DNAM-1 and CD56 in human antibody-expanded Vδ1 and Vδ2 subsets, and zoledronic-expanded (Zol) Vδ2 subsets after IL-12/18/21 activation were measured by flow cytometry. Percentages and MFI levels are shown. Data is obtained from three donors. Data is representative of three independent experiments. Zoledronic-expanded human γδ T cells from healthy donors with Vδ2^+^>80% purity were used in these experiments. Data shown as mean ± SEM. Significance was determined by one-way ANOVA with Dunnett’s multiple comparison test when comparing between control and cytokine pre-activated human γδT (A-B), two-way ANOVA with Tukey’s multiple comparison test (C) and students’ unpaired t-test (D). *** *p* <0.001 ** *p* <0.01 * *p* <0.05.

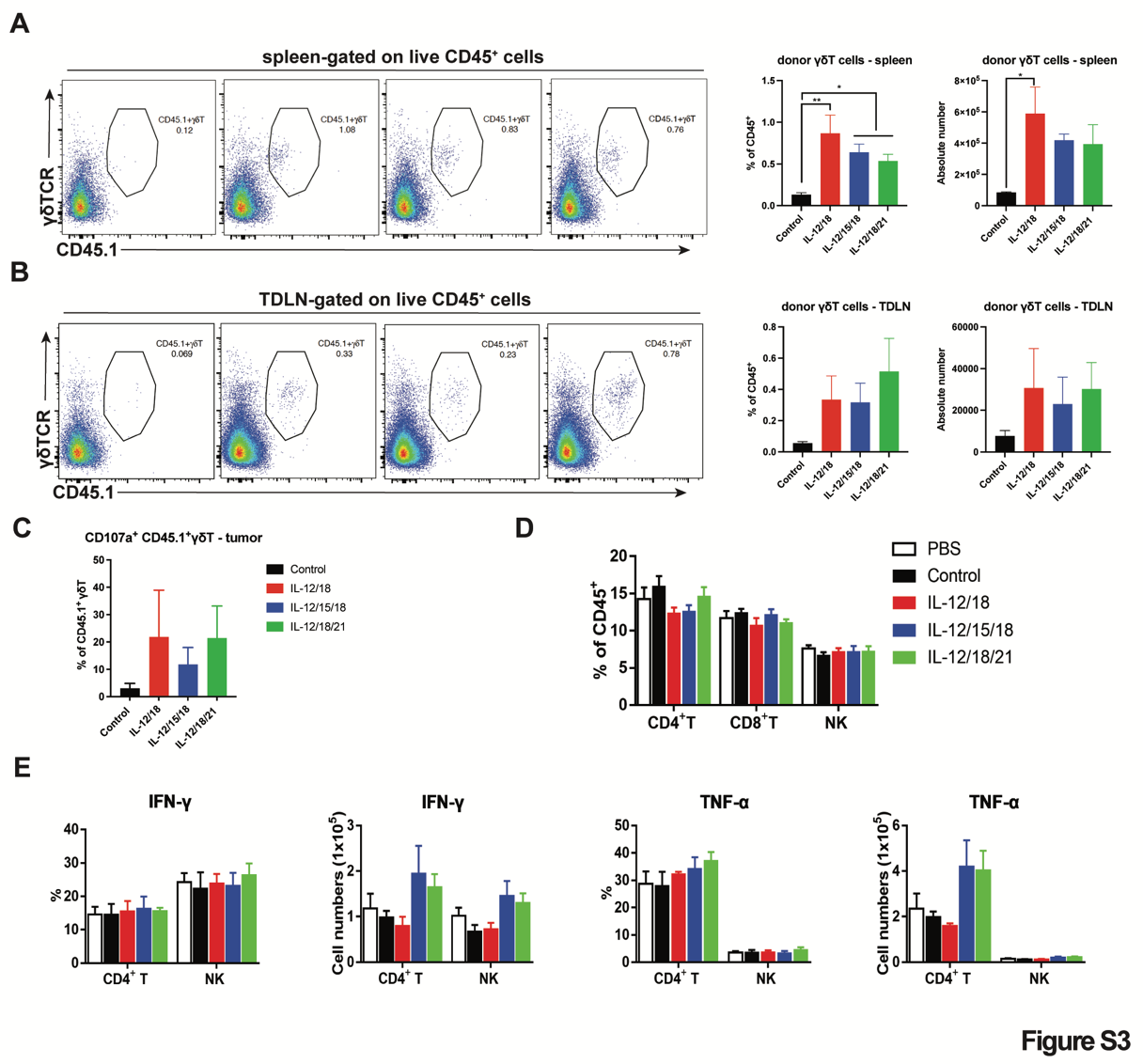

**Figure S3: Proportion of γδ T cells, CD4^+^, CD8^+^ T cells and NK cells in lymphoid organs *in vivo* after adoptive transfer to cytokine pre-activated γδ T cells.** (A-B) In B16-melanoma subcutaneous murine model, the adoptive transferred γδ T cells are detected in the spleen and tumor-draining lymph nodes (TDLN). Representative flow plots showing the CD45.1^+^ γδ T population found in spleen (top panel) and TDLN (bottom panel). Bar graphs on the right showing the percentages and cell numbers. (C) The percentage of tumor infiltrating CD107a producing donor γδT cells from subcutaneous melanoma model. The data shown is representative of 1 independent experiment with 4-5 mice per group. (D) In HCC murine model, after adoptive transfer of control γδ T cells, IL-12/18-, IL-12/15/18- and IL-12/18/21- pre-activated γδ T cells (10 x 10^6^ cells/mouse), the percentages of CD4^+^ T cells, CD8^+^ T cells and NK cells in the liver were measured by flow cytometry. (E) Percentages and cell numbers of IFN-γ-producing and TNF-α-producing CD4^+^ T cells and NK cells in the liver. The data shown are representative of 7 independent experiments. Data shown as mean ± SEM. ** *p* <0.01 * *p* <0.05, one-way ANOVA with Dunnett’s multiple comparison test.

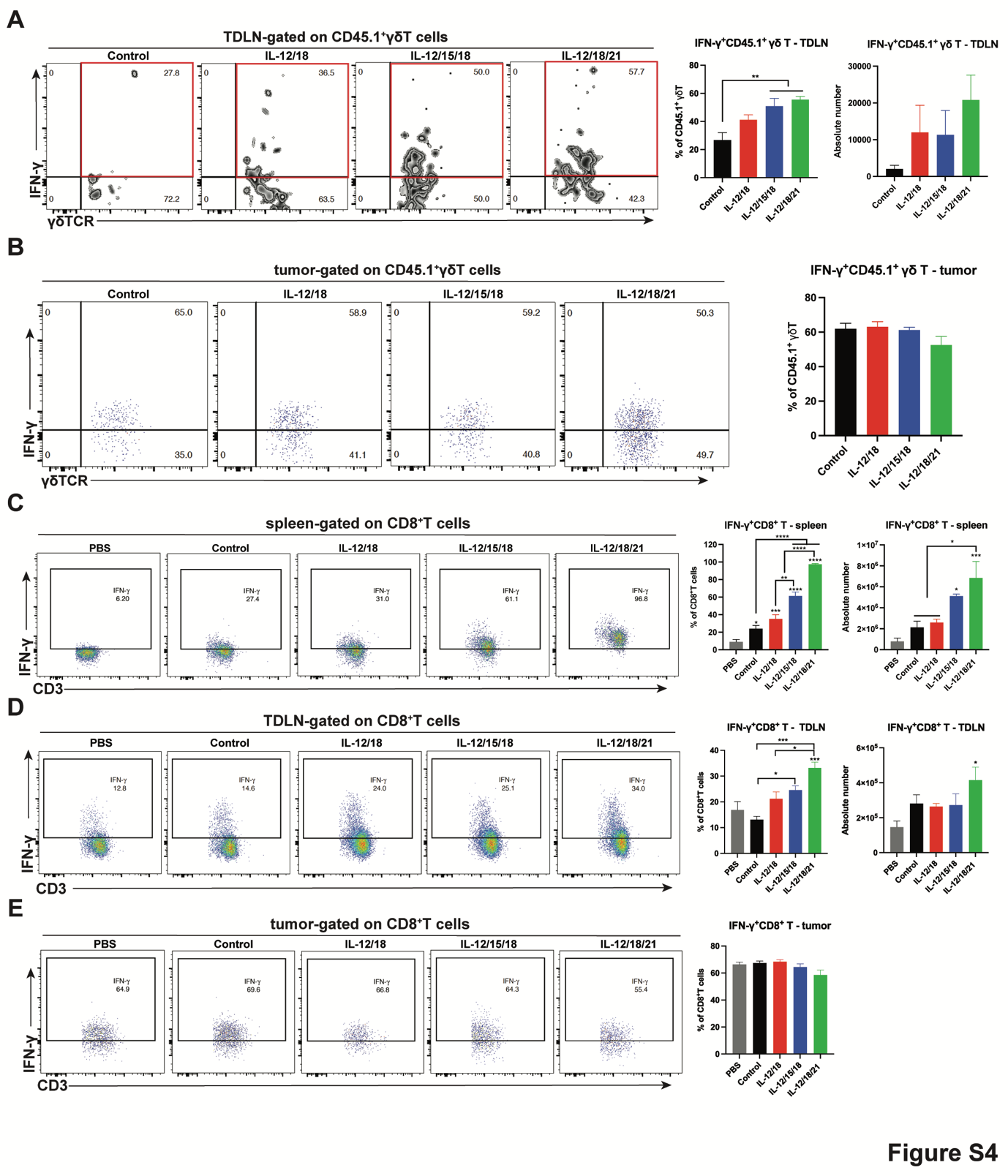

**Figure S4: IFN-γ production by adoptive transferred CD45.1^+^ γδ T cells and CD8^+^ T cells in spleen, tumor-draining lymph nodes (TDLN) and tumor CD8 in B16 melanoma murine model.** Representative contour and flow plots of IFN-γ-producing CD45.1^+^ γδ T cells in (A) TDLN and (B) tumor after adoptive transfer. The percentages and cell numbers of IFN-γ-producing CD45.1^+^ γδ T cells in TDLN and tumor are shown in the right panel. Representative flow plots of IFN-γ-producing CD8^+^ T cells in (C) spleen, (D) TDLN and (E) tumor after adoptive transfer. The percentages and cell numbers of IFN-γ-producing CD8^+^ T cells are shown in the right panel. Cells from the tumor were not counted. The data shown is representative of one independent experiment with 4-5 mice per group. Data shown as mean ± SEM. **** p<0.0001 *** p <0.001 ** p <0.01 * p <0.05, one-way ANOVA with Dunnett’s multiple comparison test.

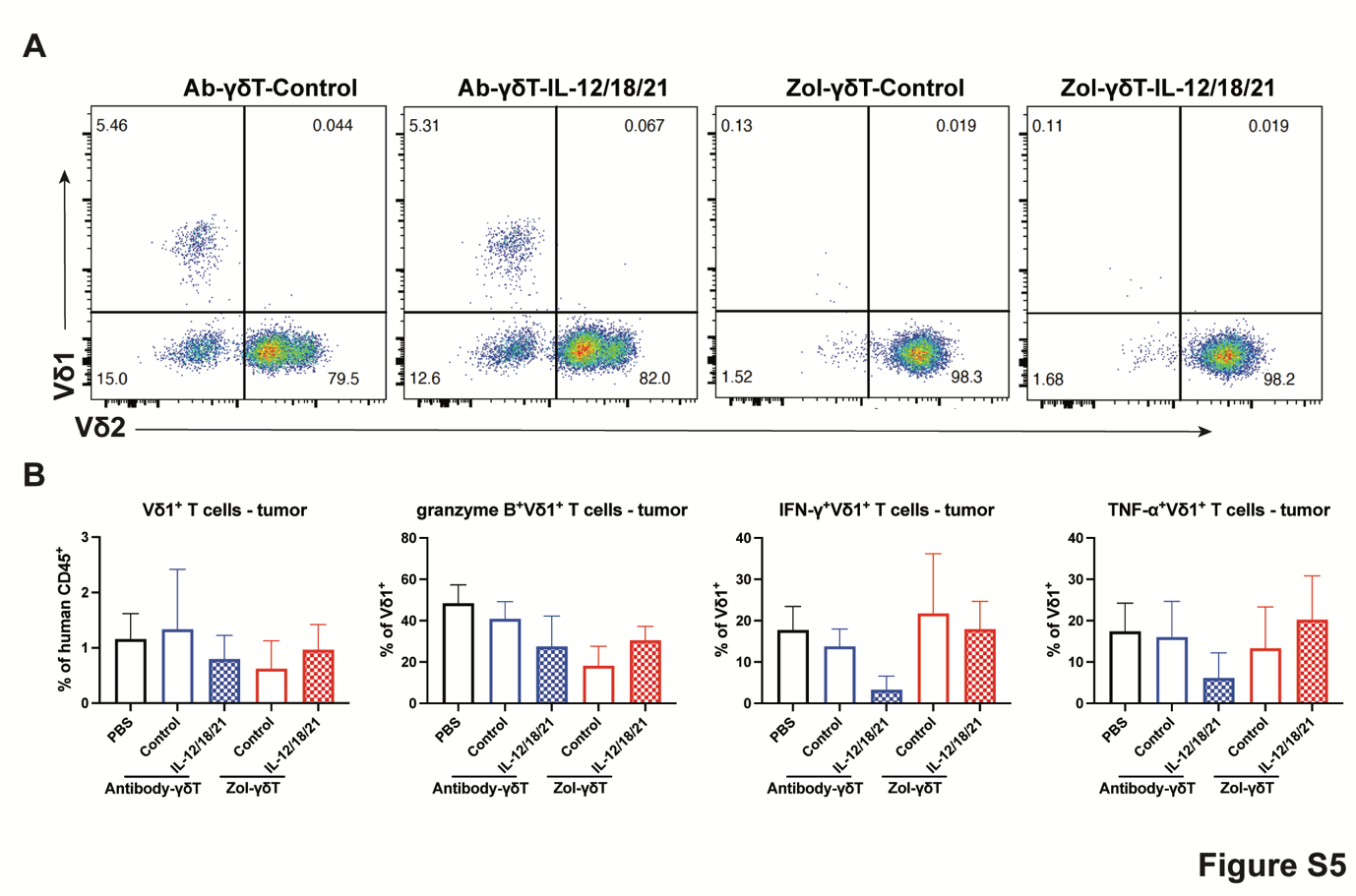

**Figure S5: Vδ1 and Vδ2 subsets after adoptive transfer of human γδ T cells in humanized murine model.** (A) The proportion of Vδ1^+^ and Vδ2^+^ T cells in tumor of mice receiving the following human γδ T cells: Antibody-γδ T-Control, Antibody-γδ T-IL-12/18/21:, Zol-γδ T-Control and Zol-γδ T-IL-12/18/21. Representative flow plots are shown, gated on human CD45^+^. (B) The percentage of Vδ1^+^ T cells in the tumor and the percentage of Granzyme B, IFN-γ and TNF-α produced by Vδ1^+^ T cells in the tumor were analyzed by flow cytometry. The data shown is representative of one independent experiment with 3-7 mice per group. Data shown as mean ± SEM. Significance was determined by one-way ANOVA with Dunnett’s multiple comparison test.

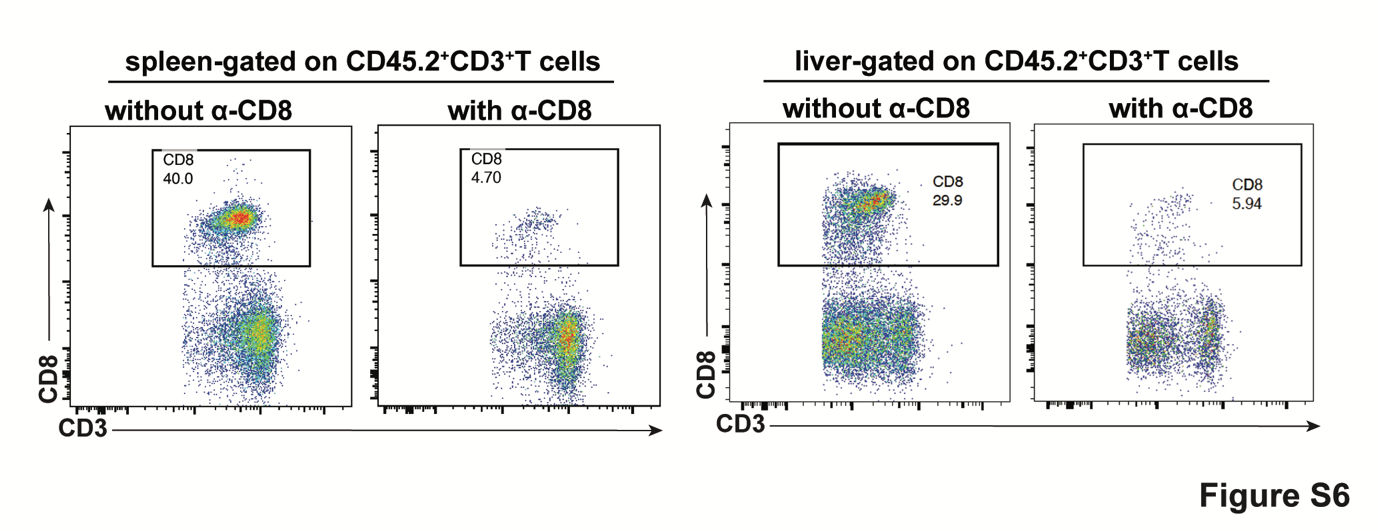

**Figure S6: Verification of CD8^+^ T cell depletion in mice.** Mice were lymphodepleted with anti-CD8 antibodies (300ug/mouse) administered via intraperitoneal route. Representative flow plots from spleen and liver showing the reduction in CD8^+^ T cells in mice receiving anti-CD8 antibody.
